# Role of the mobilome in the global dissemination of the carbapenem resistance gene *bla*_NDM_

**DOI:** 10.1101/2021.01.14.426698

**Authors:** Mislav Acman, Ruobing Wang, Lucy van Dorp, Liam P. Shaw, Qi Wang, Nina Luhmann, Yuyao Yin, Shijun Sun, Hongbin Chen, Hui Wang, Francois Balloux

## Abstract

The mobile resistance gene *bla*_NDM_ encodes the NDM enzyme capable of hydrolysing carbapenems, a class of antibiotics used to treat some of the most severe bacterial infections. *bla*_NDM_ is globally distributed across a variety of Gram-negative bacteria and is typically located within a transposon-rich genomic region common to multiple plasmids. We compiled a dataset of over 2000 bacterial genomes harbouring the *bla*_NDM_ gene including 112 new PacBio hybrid assemblies from China and developed a novel computational approach to track structural variants in bacterial genomes. We were able to correlate specific structural variants with plasmid backbones, bacterial host species and sampling locations, and identified multiple transposition events that occurred during the global dissemination of *bla*_NDM_. Our results highlight the importance of transposons in the global spread of antimicrobial resistance genes and suggest that genetic recombination, rather than mutation, was the dominant force driving the evolution of the *bla*_NDM_ genomic region.

## Introduction

The increasing burden of antimicrobial resistance (AMR) poses a major challenge to human and veterinary health. AMR can be conferred by vertically inherited point mutations or via the acquisition of horizontally transmitted non-essential ‘accessory’ genes generally located in transposons and plasmids. The *bla*_NDM_ gene encoding the NDM enzyme, a metallo-β-lactamase capable of hydrolysing most β-lactam antibiotics represents a typical example of a mobile antibiotic resistance gene (Wu et al., 2019). Compounds belonging to the carbapenem class are commonly employed to treat Gram-negative bacterial infections resistant to mainstay antibiotics and used as first-line treatment for severe infections. The global prevalence of bacteria carrying *bla*_NDM_, including carbapenem-resistant *Acinetobacter baumannii* and *Enterobacteriaceae* in hospital settings, represents a major public health concern.

The *bla*_NDM_ gene was first identified in 2008 from a *Klebsiella pneumoniae* isolated from a urinary tract infection in a Swedish patient returning from New Delhi, India (Yong et al., 2009). While *bla*_NDM_ now has a worldwide distribution, most of the earliest cases have been linked to the Indian subcontinent, suggesting this region as a likely location for the initial mobilisation event (Castanheira et al., 2011; Kumarasamy et al., 2010; Poirel, Dortet, et al., 2011; Struelens et al., 2010; Wu et al., 2019). Notably, NDM-positive *Acinetobacter baumannii* isolates have been retrospectively identified from an Indian hospital in 2005 (Jones et al., 2014), which remain the earliest observations to date. However, an NDM-positive *A. pittii* isolate was also isolated in 2006 from a Turkish patient with no history of travel outside Turkey (Roca et al., 2014).

Although no complete genome sequences are publicly available from these earliest observations, the first NDM-positive isolates from 2005 were shown to carry *bla*_NDM_ on multiple non-conjugative, but potentially mobilizable plasmid backbones (Jones et al., 2014). In addition, *bla*_NDM_ in these early isolates was positioned within a complete Tn*125* transposon with existing ISCR*27* and IS*26* insertion sequences (ISs), suggesting the possibility of complex patterns of mobility since the gene’s initial integration. Subsequent NDM-positive isolates, spanning a range of species, consistently harbour either a complete or fragmented IS*Aba125* (an IS constituting Tn*125*), which is always found immediately upstream of *bla*_NDM_ providing a promoter region for the gene transcription (Poirel, Bonnin, et al., 2011; Poirel, Dortet, et al., 2011; Toleman et al., 2012; Wu et al., 2019). The presence of IS*Aba125*, in some form, in all NDM-positive isolates to date, as well as the majority of the early observations being in *A. baumannii*, has led to Tn*125* being proposed as the likely transposon responsible for the initial mobilization of *bla*_NDM_, and *A. baumannii* as the ancestral host.

In addition, the NDM enzyme itself has been described as of possible chimeric origin (Partridge & Iredell, 2012; Toleman et al., 2012), with the first six amino acids in NDM matching to those in *aphA6*, a gene providing aminoglycoside resistance and also flanked by IS*Aba125*. It is presumed that ISCR*27*, an IS which uses a rolling-circle (RC) transposition mechanism (Ilyina, 2012; Toleman et al., 2006), initially mobilized a progenitor of *bla*_NDM_ in *Xanthomonas sp*. and placed it downstream of IS*Aba125* (Partridge & Iredell, 2012; Poirel et al., 2012; Sekizuka et al., 2011; M A Toleman, Spencer, Jones, & Walsh, 2012). The NDM enzyme itself displays some polymorphism, with at least 29 distinct sequence variants having been described to date. The most prevalent of these variants is the first to have been characterised, and is denoted NDM-1 (Basu, 2020). Different NDM variants are mostly distinguished by a single amino-acid substitution, with the exception of NDM-18 which carries a tandem repeat of five amino acids. None of the observed substitutions occur in the active site and the functional impact of each of these substitutions remains under debate (Wu et al., 2019).

At present, NDM resistance is globally distributed and represents a major concern in healthcare settings. The gene is found in at least 11 bacterial families and NDM-positive isolates have heterogeneous clonal backgrounds, supporting multiple independent acquisitions of *bla*_NDM_ (Wu et al., 2019). The *bla*_NDM_ gene has been observed on bacterial chromosomes (Baraniak et al., 2016; Rahman et al., 2018) but is most commonly harboured on plasmids, comprising multiple different backbones or types. Furthermore, even within the same plasmid types, *bla*_NDM_ is found in a variety of genetic contexts, often interspersed by multiple ISs and composite transposons (Partridge & Iredell, 2012; Wu et al., 2019). The immediate genetic environment of *bla*_NDM_ has been reported to vary even in isolates from the same patient (Wailan et al., 2015). It is therefore clear that the emergence and subsequent dissemination of NDM resistance, through a multitude of bacterial host species, is a dynamic and multi-layer process involving multiple mobile genetic elements – ‘the mobilome’ – which abetted the mobility of *bla*_NDM_ via a diverse set of processes, including genetic recombination, transposition, conjugation, transformation, and transfer through outer-membrane vesicles (OMVs) (Chatterjee et al., 2017; González et al., 2016; Huang et al., 2013; Lynch et al., 2016).

In this work, we reconstruct the individual roles of plasmids and ISs in the dissemination of NDM and provide a comprehensive overview of the many genetic backgrounds harbouring the *bla*_NDM_ gene. To this end, we compiled a global dataset of more than 2000 NDM-positive isolates including 112 newly generated hybrid PacBio assemblies sampled from clinical and livestock settings across China. In order to decompose the high sequence complexity of the immediate genomic contexts of *bla*_NDM_ in our large global dataset, we developed a novel alignment-based method designed to uncover all structural variations flanking *bla*_NDM_. This allowed us to pinpoint individual insertion events for subsequent assessment. Correlating specific structural variants with plasmid backbones, bacterial host genera and sampling locations, we are able to uncover transposition events underlying the global spread of *bla*_NDM_. We identify Tn*125*, Tn*3000* and IS*26* as the main contributors to *bla*_NDM_ mobility. Furthermore, we provide evidence for genetic recombination being the main force driving evolution in this region. We also identify plasmid backbones and bacterial hosts closely associated with specific sampling locations, as well as an apparent plateau in the rate of spread of *bla*_NDM_ around 2014. Our findings position plasmids as the main contributors to the local transmission of *bla*_NDM_, while transposons seem to be more influential for spread at a global scale.

## Results

### A global dataset of *bla*_NDM_ carriers

To study the genetic context and global spread of the *bla*_NDM_ resistance gene, a dataset of 2,148 bacterial genomes (2,166 contigs) carrying at least one copy of *bla*_NDM_ were compiled from multiple sources (Figure 1). These include: 795 bacterial genomes assembled using short read *de novo* assembly methods; 113 bacterial genomes using hybrid PacBio-Illumina *de novo* assembly; and 1,240 RefSeq assemblies (See Methods, Supplementary Table 1). Of the included *de novo* hybrid assemblies, 112 were newly generated for this study isolated from 87 hospitalized patients across China and 25 livestock farms. Overall, the dataset includes NDM-positive genomes sampled across 67 states (Figure 1A). The majority of isolates were collected in East and South East Asian countries with mainland China representing the predominant source of origin (*n*=668). A wide range of bacterial species were represented with *Klebsiella* and *Escherichia* the primary genera each contributing 899 and 667 genomes, respectively (Figure 1B; Supplementary Data 1).

**Figure 1.**
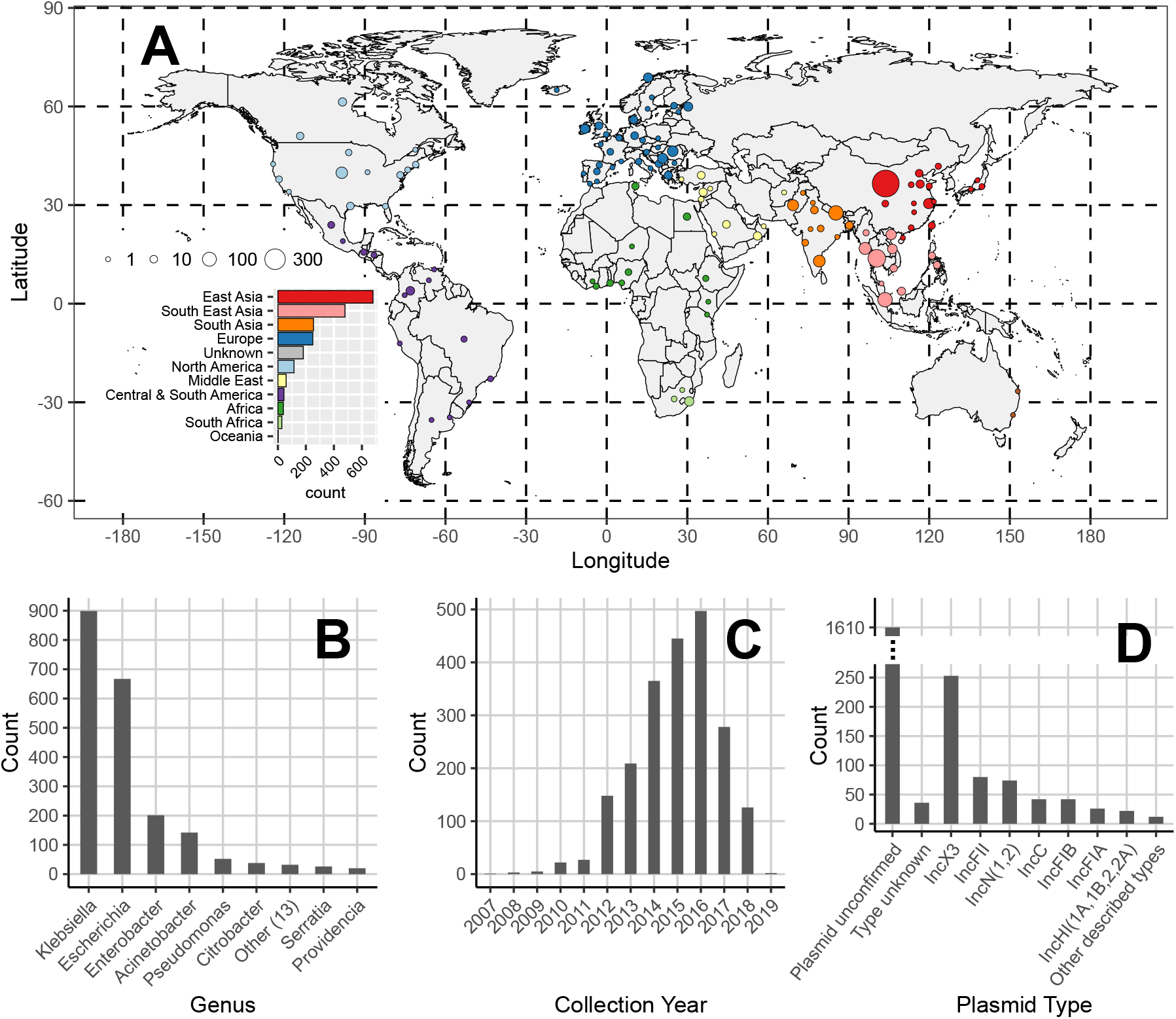
Composition of the global dataset of 2,148 NDM-positive samples. (**A**) Geographic distribution of NDM-positive assemblies. Points are coloured by geographic region and their size reflects the number of samples they encompass. **(B)** Distribution of host bacterial genera of NDM-positive samples. **(C)** Distribution of sample collection years. **(D)** Identified plasmid types on contigs bearing the NDM-resistance. All uncircularized contigs with unknown plasmid type were labelled ‘plasmid unconfirmed’. On the other hand, all circularized contigs with an unknown plasmid type were still considered plasmids but labelled ‘type unknown’.

The majority of *bla*_NDM_ carriers in the global dataset were collected between 2014-2017 (74.41%, Figure 1C). However, the dataset also includes 31 genomes from 2010 and earlier. These include the *K. pneumoniae* isolate from 2008 in which *bla*_NDM_ was first characterized (Yong et al., 2009), as well as an earlier *A. baumannii* isolate from 2007 in an individual of Balkan origin in Germany (Bonnin et al., 2012; Sahl et al., 2015) (Supplementary Data 1).

A substantial number of contigs isolated from our dataset were sufficient in length to enable identification of putative plasmid backbones carrying *bla*_NDM_ (Supplementary Figure 1; See Methods). Within our filtered dataset comprising 2,142 contigs (see Methods), we identified 482 replicon types using PlasmidFinder (Carattoli et al., 2014) and 194 circularized contigs in our dataset, of which 43 did not have a known replicon type. This resulted in a total of 525 putative plasmid sequences which also comprised 96 contigs (70 circularized) from our hybrid PacBio-Illumina assemblies. Overall, 32 different plasmid replicon types were identified among *bla*_NDM_-containing plasmid sequences (Figure 1D). The most prevalent replicon type was IncX3, found in almost half (253/525, 48%) of the included sequences. Nevertheless, the notable range in plasmid backbones harbouring *bla*_NDM_ indicates a high recombination and/or transposition rate of the *bla*_NDM_ gene. At the same time, we observe some geographic structure in plasmid replicon types (Supplementary Figure 2) signalling the importance of transposon movement in the cross-continental spread of NDM-mediated resistance.

### Resolving structural variants in the *bla*_NDM_ flanking regions

To gain a detailed overview of the transposition events and different genetic backgrounds harbouring *bla*_NDM_ we developed an alignment-based approach to resolve structural variation in the genetic regions flanking *bla*_NDM_ (see Methods, Figure 2). In brief, a pairwise discontiguous Mega BLAST search (v2.10.1+) (Camacho et al., 2009; Ma et al., 2002) was applied to all *bla*_NDM_-containing contigs in order to identify all possible homologous regions between each contig pair. Only BLAST hits covering the complete *bla*_NDM_ gene were retained (Figure 2A). Next, starting from *bla*_NDM_, a gradually increasing ‘splitting threshold’ was introduced to monitor structural variants as they appeared upstream or downstream of the gene. At each step, a network is constructed connecting contigs (nodes) that share a BLAST hit with a minimum length as given by the ‘splitting threshold’ (Figure 2B). As we move upstream or downstream and further away from the gene, the network starts to split into smaller clusters each carrying contigs that share an uninterrupted stretch of homologous DNA. The splitting is visualized as a tree where branch lengths are scaled to match the position within the sequence, and the thickness and the colour intensity of the branches corresponds to the number of sequences which are homologous (Figure 2C). Given the approach uses the *bla*_NDM_ gene as an anchor, it enables comparison between BLAST hits, but also limits the comparison to either upstream or downstream flanking region and not both simultaneously.

**Figure 2.**
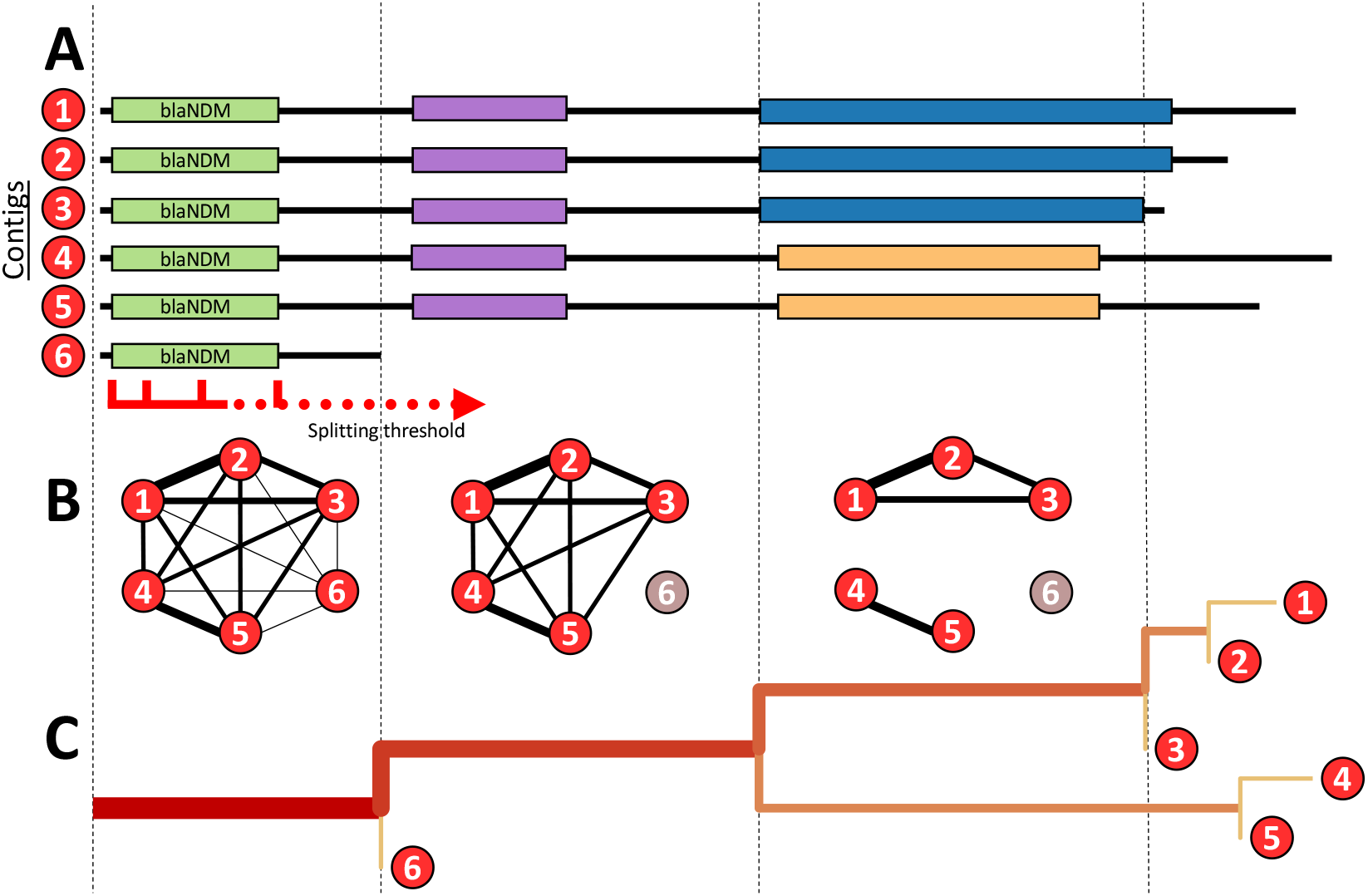
Schematic representation of the tracking algorithm splitting structural variant bacgrounds upstream or downstream of *bla*_NDM_ gene. **(A)** A pairwise BLAST search is performed on all NDM-positive contigs. Starting from *bla*_NDM_ and continuing downstream or upstream, the inspected region is gradually increased using the ‘splitting threshold’. **(B)** At each step, a graph is constructed connecting contigs (nodes) that share a BLAST hit with a minimum length as given by the ‘splitting threshold’. Contigs which have the same structural variant at the certain position of the threshold belong to the same graph component, while the short contigs are singled out. **(C)** The splitting is visualized as a tree where branch lengths are scaled to match the position within the sequence, and the thickness and the colour intensity of the branches correspond to the number of sequences carrying the homology.

The flanking region upstream of *bla*_NDM_ breaks down rather quickly: within a few hundred base pairs of the *bla*_NDM_ start codon, the upstream flanking region splits into multiple structural variants, none of which dominates the contig pool (Supplementary Figure 3). For instance, 99 different structural variants were identified only 1200 bp from the *bla*_NDM_ start codon. This high variation in genome structure could be attributed to the many genetic backgrounds in which *bla*_NDM_ is found as well as frequent genome rearrangements (Supplementary Figures 3). The significance of the latter is also reflected by the number of fragments and complete insertion sequences present in the region, including IS*Aba125* (132), IS*5* (385), IS*3000* (88), IS*Kpn14* (44), and IS*Ec33* (72), as well as almost half the contigs (1,003, 46.93%) being excluded from the analysis for having too short an upstream flank (Supplementary Figure 3). The transposition hotspot upstream of *bla*_NDM_ possibly hinders sequencing and genome assembly efforts and enhances the presence of these short contig flanks. In agreement with previous work (Poirel, Bonnin, et al., 2011; Poirel, Dortet, et al., 2011; Toleman et al., 2012; Wu et al., 2019), more than 95% of sufficiently long contigs include a ~75 bp fraction of IS*Aba125*, supporting the notion of Tn*125* as an ancestral transposon of the *bla*_NDM_ gene (Supplementary Figures 3 and 4).

The downstream flanking region exhibits more gradual structural diversification than the upstream region, with one dominant putative ancestral background (Figure 3). As illustrated by the stem of the tree of structural variations (Supplementary Figure 5), many of the 2,142 contigs analysed contain complete sequences of the same genes: *ble* (2,047 contigs), *trpF* (1,770), *dsbD* (1,660), *cutA* (858), *groS* (673), *groL* (527). In total there are 1,229 contigs which are sufficiently long downstream of *bla*_NDM_ to harbour the full repertoire of the aforementioned genes. When the analysis is restricted to those contigs of sufficient length, 42.9% of NDM-positive contigs carry this full suite of genes downstream of *bla*_NDM_.

**Figure 3.**
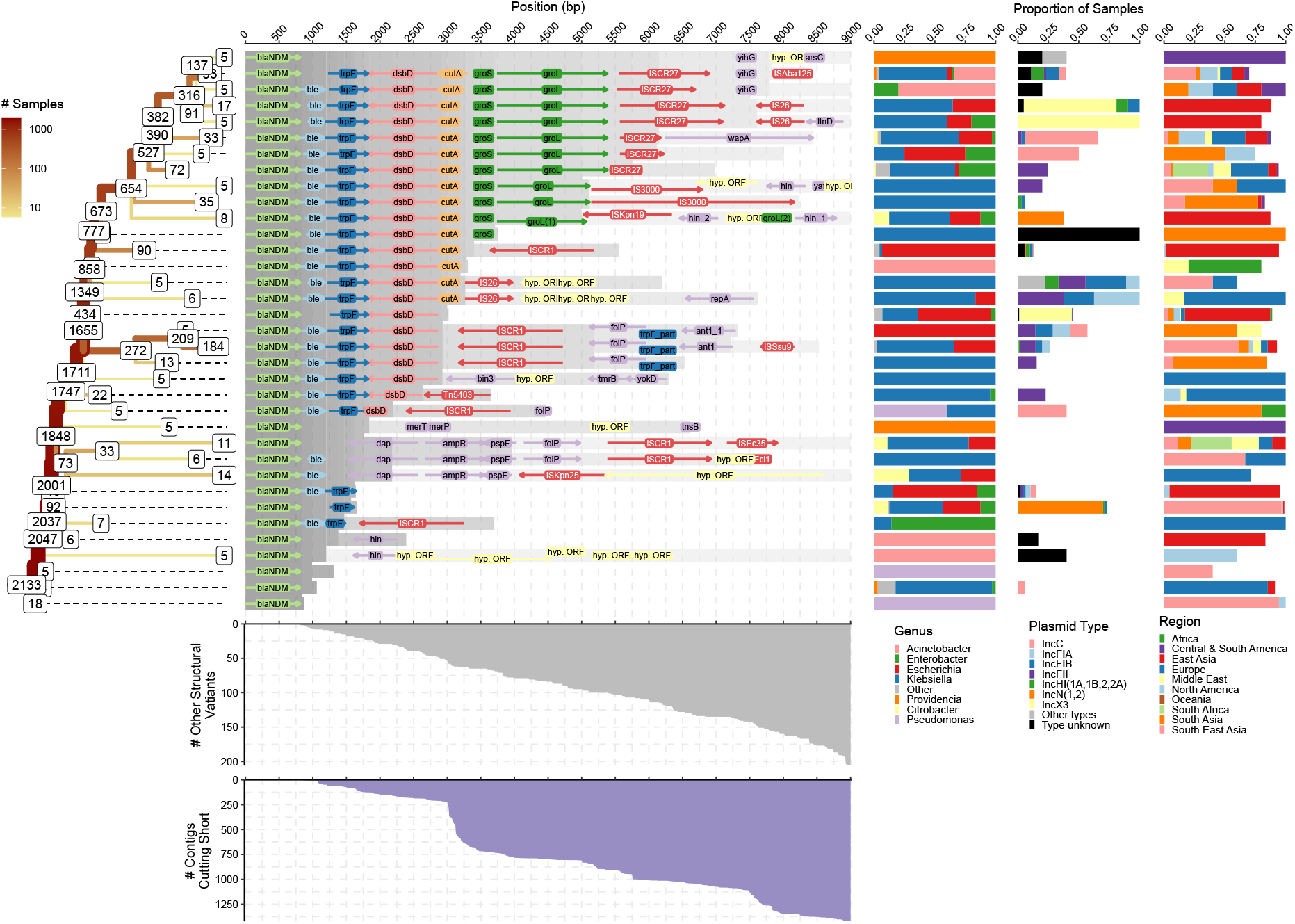
Splitting of structural variants downstream of *bla*_NDM_. The ‘splitting’ tree for the most common (i.e ≥ 5 contigs) structural variants is shown on the left-hand side. The labels on the nodes indicate the number of contigs remaining on each branch. The other contigs either belong to other structural variants or were removed due to being too short in length. The number of contigs cutting short is indicated by the area chart at the bottom. Similarly, the number of contigs belonging to less common structural variants is indicated by the upper area chart. The genome annotations of most common structural variants are shown in the middle of the figure. The homologous regions are indicated by the grey shading. Some of the structural variants and branches were intentionally cut short even though their contigs were of sufficient size. This was done in order to prevent excessive bifurcation and to make the tree easier to interpret. In particular, branches with percent change of contigs lost due to variation and shortness above 10% were truncated. The distribution of genera, plasmid types and geographical regions of samples that belong to a each of the common structural variant is shown on the right-hand side.

### Patterns of insertion events in *bla*_NDM_ flanking regions

Having reconstructed structural variation in the *bla*_NDM_ upstream (Supplementary Figure 3) and downstream (Figure 3) flanking regions, we did not observe any strong overall signal in the distribution of associated plasmid backbones, bacterial genera and sampling locations. However, closer examination of structural variants common to sufficiently large pools of isolates allow distinct observations to be made. These more specific observations appear to correlate to the events underlying the spread of *bla*_NDM_. For instance, IS*3000* is found in 88 and 35 contigs on the upstream and downstream flanking regions respectively, almost exclusively in *Klebsiella* host species and often on IncF plasmids (Figure 3 and Supplementary Figure 3). Thus, as previously suggested by Campos et al., Tn*3000* likely re-mobilized *bla*_NDM_, following the fossilization of Tn*125* (Campos et al., 2015); our analysis suggests the secondary mobilization primarily happened in *Klebsiella* species.

Some structural variants appear geographically linked e.g., IS5 is predominantly found upstream of *bla*_NDM_ on IncX3 plasmids from East Asia (Supplementary Figure 3), with none of these plasmids with IS5 having a matching element on the downstream flanking region of *bla*_NDM_ to form a full composite transposon. IS5 is known to enhance transcription of nearby promoters in *E. coli* (Schnetz & Rak, 1992) and its abundance and positioning just upstream of *bla*_NDM_ suggests it may have assumed a similar role in this case. Interestingly, the NDM-5 variant has been increasing in numbers in recent years (Supplementary Figure 6 A and B) and is mostly associated with both IncX3 plasmids (Supplementary Figure 6 C and D) and isolates from East Asia (Supplementary Figure 6 G and H). Thus, an increasing abundance of NDM-5 could be due to the aforementioned enhanced transcription caused by the proximity to IS5. Other structural variants are observed across many global regions e.g., the *wapA* gene is found truncating ISCR27 downstream on IncC plasmids (Figure 3).

One of the most commonly found transposable elements in the flanking regions (~30% prevalence) is an ISCR1-like transposase (IS91 family transposase), hereafter referred to as ISCR1, coupled with the *folP* gene (Figure 3, Supplementary Figure 5). This configuration is found at various positions downstream of *bla*_NDM_ and often associated to IncF plasmids identified in *Escherichia* and *Klebsiella* species. In most cases, the orientation of ISCR1 should prevent this element from mobilizing *bla*_NDM_ (Ilyina, 2012), so it appears its role is to disrupt the surrounding IS elements and transposons. Interestingly, ISCR1s are mainly found in complex class 1 integrons (Ilyina, 2012), however, not many annotated integrase genes are located within the vicinity of *bla*_NDM_. In fact, only 11 contigs were found to have an integrase <50 Kb away from *bla*_NDM_ and none showed any consistency in how the integrase is placed with respect to *bla*_NDM_. This may suggest integrons play at most a minor role in the dissemination of *bla*_NDM_.

Another notable ISCR element is ISCR27 which is consistently found immediately downstream of the *groL* gene (Figure 3, Supplementary Figure 5). The complete ISCR27 sequence is carried by 316 contigs, with another 211 contigs containing a fragmentary sequence. ISCR27 is found at high prevalence with 30.1% of sufficiently long contigs harbouring this element. Contrary to its ISCR1-like relative, ISCR27 is correctly oriented to mobilize *bla*_NDM_ as is presumed to have happened during the initial mobilization of the progenitor of *bla*_NDM_ (Toleman et al., 2012). However, we find no evidence of subsequent ISCR27 mobility. The origin of rolling-circle replication of ISCR27 (*oriIS*; GCGGTTGAACTTCCTATACC) is located 236 bp downstream of the ISCR27 transposase stop codon. The region downstream of this stop codon in all structural variants bearing a complete ISCR27 is highly conserved for at least 750 bp (Figure 3, Supplementary Figure 5). This suggests a reasonably conserved genetic background surrounding ISCR27 as *bla*_NDM_ has been disseminated.

Surprisingly, only 58 contigs carried a complete IS*Aba125* downstream of *bla*_NDM_, of which 53 carried an IS*Aba125* sequence in proximity (<7886 bp) to the *bla*_NDM_ start codon. These account for a minority (7.4%) of isolates when sufficiently long contigs are considered. Forty-five of these contigs contained a complete IS*Aba125* both upstream and downstream of *bla*_NDM_ thus forming a complete Tn*125* transposon. Even though the diversity of bacterial genera carrying IS*Aba125* upstream is substantial (Supplementary Figure 3), the less preserved downstream IS*Aba125* sequence is mostly found in the genera *Acinetobacter* and *Klebsiella* (Figure 3). This supports the initial dissemination of *bla*_NDM_ by Tn*125* to other plasmid backbones predominately being mediated by these two genera, after which the transposon was disrupted by other rearrangements.

We note that more than 500 contigs were truncated around 3000 bp downstream of *bla*_NDM_ (Figure 3). To investigate the reasons behind this distinct cut-off point, we used 447 raw short-read sequencing samples from our dataset (originally downloaded from SRA, see Methods) with contigs that carry *bla*_NDM_ longer than 3000bp (Supplementary Table 1). We compared the normalized number of reads with overhangs mapping to the end of contigs ending 3000-3200 bp and longer contigs, ending >3200 bp downstream of *bla*_NDM_ (Supplementary Figure 7A). On average, the normalized number of overhangs is two times higher in shorter contigs, which indicates that a particular genetic region mapped by the overhanging reads is often present in more than one copy. Moreover, when mapped back to the assembled contigs, the overhanging reads of shorter contigs are found on average on three different contigs (>1000 bp) – twice as many as observed for longer contigs (Supplementary Figure 7B). The presence of these overhanging reads on multiple contigs may point to within-isolate transposition/rearrangement events between plasmids and/or bacterial genomes which seem to localise around this region.

The shorter contigs (3000-3200 bp) are found across genera of *Enterobacteriaceae* including *Escherichia*, *Klebsiella*, *Enterobacter*, *Citrobacter*, *Leclercia* and *Lelliottia*. What is more, the overhanging reads of shorter contigs almost exclusively match the left inverted repeat (IRL) of IS*26* sequence. In fact, over one third (157; 35.1%) of all analysed contigs’ overhanging reads correspond to IS*26* IRL. IS*26*, although often found in two adjacent copies forming a seemingly composite transposon, is a so-called pseudo-composite (or pseudo-compound) transposon (Harmer et al., 2020). In contrast to composite transposons, a fraction of DNA flanked by the two IS*26* is mobilized either via cointegrate formation or in the form of a translocatable unit (TU), which consists of a single IS*26* element and a mobilized fraction of DNA, and inserts preferentially next to another IS*26* (Harmer et al., 2014, 2020). Interestingly, no IS*26* sequences were found upstream within contigs whose downstream overhanging reads match to IS*26*. Assembly procedures are known to struggle with allele duplications which may explain the lack of IS*26* sequences upstream and the surge of truncated contigs (Sohn & Nam, 2018). Nevertheless, the results above suggest an active within-isolate movement of *bla*_NDM_ via IS*26* across *Enterobacteriaceae*.

In total, we identified 208 putative composite transposons (i.e., stretches of DNA flanked by at least two ISs enclosing *bla*_NDM_ <30 Kb apart) in 181 contigs. These comprised 18 different types with the five most frequent being: IS*26* (62 instances), IS*Aba125* (forming Tn*125*; 55 instances), IS*3000* (forming Tn*3000*; 52), IS*15* (13), IS*6100* (7). Interestingly, there are 38 cases where >2 of the same IS flank *bla*_NDM_. These are mostly IS*26* (23). Also, only 137 of the 208 putative transposons identified contained both complete flanking ISs, while others had at least one IS partially truncated. Importantly, IS*26*, IS*6100* and IS*15*, a known variant of IS*26*, are phylogenetically related with all three falling into clade I of the IS*6* family of insertion sequences whose members are known to mobilize via cointegrate formation, as discussed above (Harmer & Hall, 2019). The IS26s we identify are found at different positions in the alignment, usually between 10-20 Kb apart, while other ISs are, for the most part, found at a fixed position around *bla*_NDM_. This indicates increased activity and multiple independent acquisitions of IS*26*. As expected, the transposons we identify are found on various plasmid backbones (Supplementary Figure 8C). However, some trends can be identified in the distribution of associated bacterial genera and geographic region of sampling (Supplementary Figure 8A and B). In particular, Tn*3000* is almost exclusively found in *Klebsiella* species and Tn*125* predominantly in *Acinetobacter* and *Klebsiell*a, while IS*26* are found in *Escherichia* and *Klebsiella*. In spite of these elements being present across the globe, some geographic structure is apparent. For example, IS*26* appears to dominate in East Asia while Tn*3000* tends to occur in South Asia. Overall, the distributions of various structural variants and transposons with respect to plasmid replicon types and bacterial hosts suggest that most rearrangements in the *bla*_NDM_ flanking regions happened within *Escherichia* and *Klebsiella* species where IS26, Tn*125* and Tn*3000* are the main contributors to *bla*_NDM_ mobility.

### Mutations accumulated in *bla*_NDM_ transposons provide only weak evolutionary signal

To further investigate the dynamics of spread of the *bla*_NDM_ gene, regression analyses and Bayesian molecular tip-dating (implemented in BEAST2 v2.6.0) (R. Bouckaert et al., 2019) were performed on full alignments of Tn*125* (45 contigs) and Tn*3000* (29 contigs) (Supplementary Figure 9). SNPs within each alignment were identified using a consensus sequence approach (see Methods). Few SNPs are observed in the alignments of Tn*125* (56 SNPs) and Tn*3000* (14) (Supplementary Figure 9A and B). In fact, a general observation was that relatively few SNPs are found in alignments of any stretch of homologous sequence flanking *bla*_NDM_ relative to the number of structural variants. For instance, only 80 SNPs are present in the 2,570 bp alignment of 1,711 contigs harbouring *bla*_NDM_, *ble*, *trpF*, and *dsbD* genes, while more than 50 different structural variations are found over the same distance downstream of the *bla*_NDM_ start codon. Going downstream, the number of structural variants increases while the number of newly accumulating SNPs plateaus, as fewer samples are available and the genetic background diversifies.

This restricted genetic diversity of the two transposon alignments results in only a weak temporal signal (see Methods and Supplementary Figure 9A and B). While results should therefore be interpreted with appropriate caution, we proceeded with Bayesian molecular tip-dating analyses to assess the relative timing of transposition events involving Tn*125* and Tn*3000* (see Methods). All models converged well, though we note that both marginal distributions of the most common recent ancestor (tMRCA) of Tn*125* and Tn*3000* (Supplementary Figure 9C and D) overlap with the marginal distributions of the corresponding model priors (i.e., BEAST2 runs without SNP data provided) (Supplementary Figure 9D) which is a likely consequence of the lack of genetic diversity. Nevertheless, the tMRCA estimates of Tn*125* and Tn*3000* shift from the expectation under the priors. In particular, the Tn*3000* marginal distribution points to a later date indicating that the tMRCA of Tn*3000* carrying *bla*_NDM_ gene emerged after mid-2008, but still before the earliest sampling date at the end of 2011 (Supplementary Figure 9C). In contrast, the marginal distribution of the Tn*125* tMRCA shifts to an earlier date, suggesting this transposon mobilized *bla*_NDM_ before 2009 and after 2004. This tMRCA distribution also includes the dates of the earliest reported Tn*125*-*bla*_NDM_-positive isolates from 2005 (Jones et al., 2014) which gives some credibility to these results.

The indications from molecular tip-dating fall into a wider narrative where *bla*_NDM_ spread was initially driven by Tn*125* mobilization before subsequent transposition by Tn*3000*, and others. However, the sparsity of SNPs within the alignments, the weak temporal signal and the abundance of structural variants, plasmid backbones, transposons and ISs argue in favour of genetic recombination, rather than *de novo* mutation, as the dominant mechanism driving evolutionary change in the genetic region flanking *bla*_NDM_ gene.

### Correlates with the global dissemination of *bla*_NDM_

The earliest samples in our dataset span the years 2007 to 2010 and comprise 31 *bla*_NDM_-positive genomes already encompassing nine bacterial species, 13 countries, and three continents (23 confirmed clinical samples and 8 of unknown origin from Asia, North America and Europe). Even though the exact time of emergence remains an open question, such a wide host and geographic distribution, even in the earliest available samples, illustrates the extraordinarily high mobility of *bla*_NDM_. To track the spread of *bla*_NDM_ we estimated diversity over time for several categorizations of *bla*_NDM_-positive samples (Supplementary Figure 11, see Methods). In particular, for each year, the diversity was estimated among samples’ country of collection, associated bacterial genera, replicon types (i.e., plasmid backbones), SNP counts within 5000 bp alignment, and structural variants at positions 3000 bp and 5000 bp downstream of the *bla*_NDM_ gene. Shannon entropy was used as a measure of diversity and bootstrapping implemented to provide confidence intervals around the entropy estimates. A strong sampling bias is present among isolates from the same NCBI BioProject (Supplementary Figure 10). To account for this, we weighted contigs during bootstrapping based on their BioProject affiliation (see Methods).

The change in diversity of the countries associated to *bla*_NDM_-positive isolates was used to approximate the broad patterns of global dissemination of NDM resistance (Supplementary Figure 11A). The diversity of sampling countries through time plateaued between 2013-2015. In light of the earliest reports of NDM-positive samples in 2005, this indicates that it took eight to eleven years for NDM resistance to spread globally and is consistent with our estimates based on phylogenetic tip-dating (Supplementary Figure 9C). Furthermore, the change in the diversity of countries associated to *bla*_NDM_-positive genomes was found to be positively correlated with all other considered categories (Supplementary Figure 12) suggesting it holds information which can be leveraged to reconstruct dissemination trends. The weakest correlation with the widest confidence interval was found between the number of SNPs in the alignment and the diversity of countries of sample origin (ρ = 0.407 [0.119-0.753]), followed by the bacterial genera (ρ = 0.5 [0.217-0.7]), then structural variants at 3000bp downstream of *bla*_NDM_ (ρ = 0.533 [0.217-0.717], and 5000bp downstream (ρ = 0.683 [0.433-0.85]). Despite the overlap of confidence intervals, this ordering again highlights the importance of genetic rearrangements and transposition in the evolution of this genetic region.

The strongest correlation was found between the diversity of countries with NDM-positive isolates and the replicon types of associated plasmid backbones (ρ = 0.7 [0.467-0.883]) supporting a strong dependence between the two (Supplementary Figure 12B). To further investigate this relationship, we assessed the correlation between genetic and geographical distance between pairs of contigs as a function of the distance downstream of *bla*_NDM_ gene (Figure 4, see Methods). Starting from *bla*_NDM_ and moving downstream, we gradually extended the region over which genetic distances were estimated. At each step, we estimated the correlation between genetic and geographic distance.

**Figure 4.**
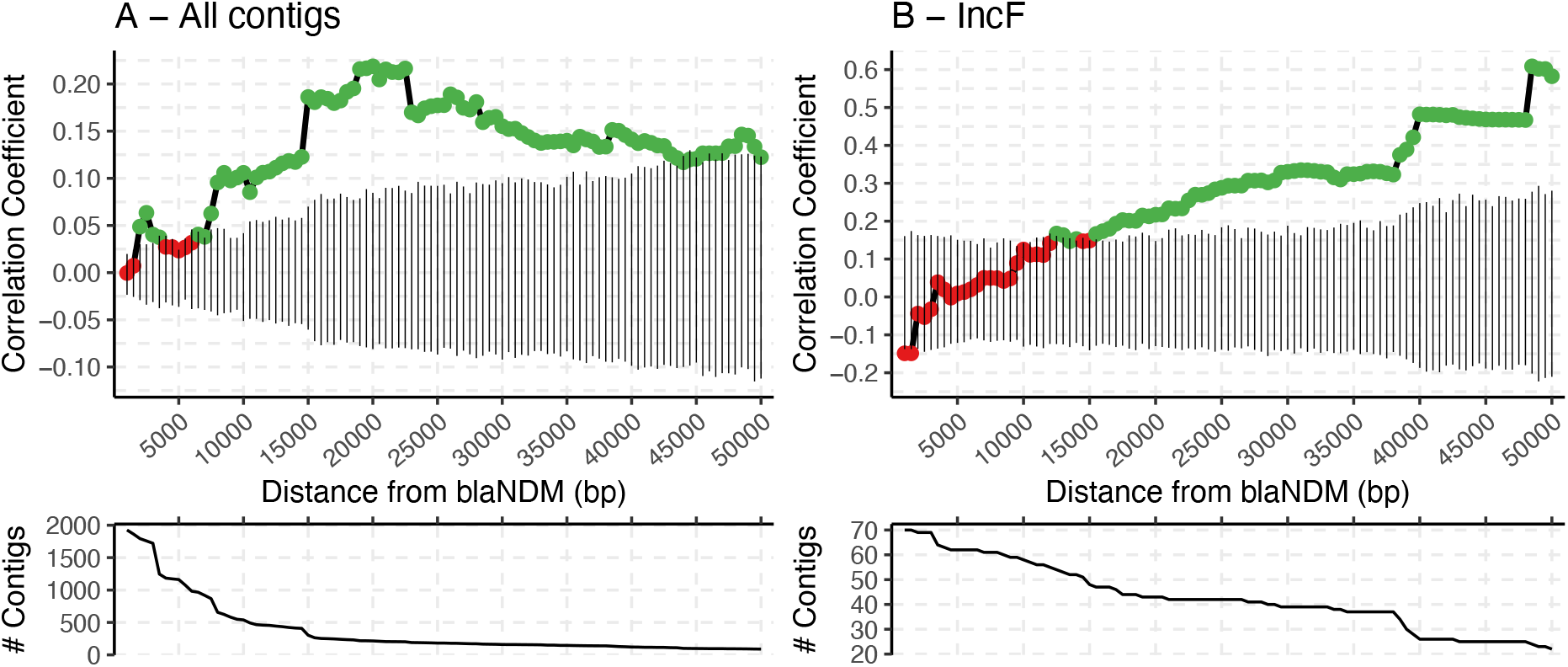
The spearman correlation estimates between genetic and geographic distance of NDM-positive contigs as the DNA sequence upon which the genetic distance is measured is increased downstream of *bla*_NDM_ gene. The exact Jaccard index, an alignment-free metric, was used as a measure of genetic distance. Geographic distance between samples was estimated by the *geodist* (v0.0.6) R package using sampling coordinates or sampling country centroids if the former had not been provided. The analysis was performed on all contigs in the dataset that carry the *bla*_NDM_ gene (**A**) and the ones with confirmed IncF replicon type (**B**). In both cases, the genetic and geographic distance was measured between all pairs of contigs from a different BioProject which yielded two distance matrices: genetic and geographic. The Spearman correlation was then estimated between the two matrices and its significance evaluated using Mantel (randomization) test. Significant Spearman correlations (p-value <0.05) are indicated with green points and non-significant correlations with the red point, while the black vertical lines provide the 95% confidence interval of 1,000 Mantel test permutations. The genetic distance matrix and subsequent Spearman correlation were estimated multiple times by increasing the assessed DNA sequence starting from *bla*_NDM_ gene and continuing downstream. The two plots below the correlation graphs indicate the number of contigs used in the correlation analysis as the assessed DNA sequence is increased. See Supplementary Figure 12 for correlation analysis on IncX3 and IncN plasmids.

Considering all contig sequences, a gradual increase in correlation between genetic and geographic distance was observed as more of the sequence downstream of *bla*_NDM_ was included (Figure 4A). The same trend is observed in an isolated case of “broad-range” IncF plasmids which have a wide geographical distribution (Figure 4B, Supplementary Figure 2). However, no significant or sufficiently long consecutive correlations were found among IncX3 and IncN plasmids (Supplementary Figure 13) likely due to the lack of longer plasmid sequences and more restricted mean geographic distance between pairs of plasmids; both replicon types are mostly found in China and India respectively (Supplementary Figure 2).

Nevertheless, considering *bla*_NDM_ is predominantly carried by plasmids (Wu et al., 2019), the trend identified in Figure 4 suggests that plasmids carrying *bla*_NDM_ are geographically structured. Gene dissemination is a fundamentally spatial process. Despite being theoretically mobile, in practice most plasmids may be both strongly host-constrained (Acman et al., 2020) and associated with particular locations or environmental niches (Shaw et al., 2020). All in all, this could be hinting at the existence of plasmid niches: settings to which particular plasmids are more adapted.

## Discussion

Increasing levels of antimicrobial resistance in bacterial pathogens pose a major global health challenge, with resistance to carbapenems a particularly concerning example. Understanding the main mechanisms by which antibiotic resistance elements are disseminated is fundamental to our understanding of the spread of AMR, and new methods are required to fully reconstruct the forces underlying the dynamic mobilome common to many resistance elements. Here, we have compiled a global dataset of 2,148 bacterial genomes carrying *bla*_NDM_, including 112 new hybrid assemblies from Chinese hospitals, to provide a comprehensive overview of the different genetic backgrounds harbouring this resistance element and to gain insight into its mobility. In order to do this, we developed a new alignment-based method to resolve the complex structural variations flanking this major antibiotic resistance element.

Our results, summarized in Figure 3, highlight the vast diversity of genetic backgrounds and plasmids harbouring *bla*_NDM_ and the predisposition of this region for genetic reshuffling. Moreover, we detected a markedly low SNP prevalence and weak temporal signal, which points to the importance of genetic recombination and transposition in driving the evolution of this region. In addition, we identified 18 different putative transposons within our dataset, of which Tn*125*, Tn*3000* and IS26 flanked pseudo-composite transposon are predominant and represent the major contributors to plasmid jumps of *bla*_NDM_. IS*26* seems particularly promiscuous; it is often found inserted at various positions around *bla*_NDM_ and with some indication of within-isolate activity. IS*26* is known for its increased activity and rearrangement of plasmids in clinical isolates (S. He et al., 2015) and has been observed to drive within-plasmid heterogeneity even in a single *E. coli* isolate (D. D. He et al., 2019). Thus, it is a likely candidate driving *bla*_NDM_ gene acquisition and extensive rearrangements found within *bla*_NDM_ region. Furthermore, IS*5* was often and uniquely found immediately upstream of *bla*_NDM_ and its peculiar positioning could foreshadow its role in increased transcription of the gene (Schnetz & Rak, 1992). Little to no evidence was found for the involvement of integrons and RC transposition of ISCR elements in spreading of *bla*_NDM_. In fact, ISCR1 alongside other ISs, was mainly found disrupting the *bla*_NDM_ region.

By assessing the change in entropy of countries where *bla*_NDM_-positive isolates have been sequenced over time, we traced the patterns underlying the spread of NDM resistance. Our assessment of diversity suggests that, following a rapid dissemination, the spread of *bla*_NDM_ may have reached a plateau between 2013-2015, with *bla*_NDM_ reaching a global prevalence 8-11 years after 2005. Such a rapid spread has also been suggested for other significant mobile resistance genes: the *mcr-1* gene, mediating colistin resistance, is also estimated to have reached global prevalence within a decade (R. Wang, Van Dorp, et al., 2018). The extent to which this model of ‘rapid spread’ applies to other transposon-borne resistance elements remains to be determined.

We found a strong positive correlation between genetic distances between plasmid backbones bearing *bla*_NDM_ and the geographic location in which they were sampled, suggesting the existence of a constraint on plasmid spread i.e. plasmid niches. We presume plasmid niches exist thanks to local evolutionary pressures for which particular plasmid backbones are optimized. Country boundaries limiting population movement, region-specific outbursts of antibiotic usage and narrow host range of the majority of bacterial plasmids (Acman et al., 2020) all likely contribute to a restricted geographical range. Thus, an introduction of another plasmid into a foreign plasmid niche may lead to plasmid loss or fast adaptation by, for instance, acquisition of resistance and other accessory elements. This hypothetical scenario also provides an opportunity for resistance to spread by transposition or recombination, by which a new resistance gene is able to enter another plasmid niche. In the case of *bla*_NDM_, this would also imply that after the initial introduction of *bla*_NDM_ to a geographic region, dissemination and persistence of the gene could proceed idiosyncratically - selection for carbapenem resistance being just one of many selective pressures acting on plasmid diversity.

The importance of transposon movement has been previously demonstrated by our work on plasmid networks (Acman et al., 2020), as well as several papers promoting a Russian-doll model of resistance mobility (Sheppard et al., 2016; R. Wang, Van Dorp, et al., 2018). In light of our results, we suggest a conceptual framework of resistance gene dissemination where plasmid mobility is for the most part restricted. Although plasmids can facilitate rapid spread within species and geographical regions, the momentum of resistance dissemination is primarily reliant on between-plasmid transposon jumps and genetic recombination.

## Methods

### Compiling the dataset of NDM sequences

We compiled a global dataset of 2,148 bacterial genomes carrying the *bla*_NDM_ gene from several publicly available databases. The vast majority of bacterial isolates were collected from patients (1,501), while 308 are of animal origin (184 from chickens, 51 from other birds and 47 from flies), 244 are of an unknown origin, and 95 are environmental samples (of which 36 are isolated from hospital environments). 1239 and 275 fully assembled genomes were downloaded from NCBI Reference Sequence Database (RefSeq; accessed on 23^rd^ of May 2019) (O’Leary et al., 2016; Pruitt et al., 2007) and EnteroBase (Zhou et al., 2020) respectively. The EnteroBase repository was screened using BlastFrost (v1.0.0) (Luhmann et al., 2020) allowing for one mismatch. In addition, we used the Bitsliced Genomic Signature Index (BIGSI) tool (Bradley et al., 2019) to identify all Sequence Read Archive (SRA) unassembled reads which carry the *bla*_NDM_ gene. At the time of writing, a publicly available BIGSI demo did not include sequencing datasets from after December 2016. Therefore, we manually indexed and screened an additional 355,375 SRA bacterial sequencing datasets starting from January 2017 to January 2019. We required the presence of 95% of *bla*_NDM_-1 *k*-mers to identify NDM-positive samples from raw SRA reads. This led to the inclusion of 522 isolates from reads downloaded from the SRA repository. Furthermore, we generated 112 new NDM-positive genomes using paired-end Illumina (Illumina HiSeq 2500) and PacBio (PacBio RS II) sequencing of isolates from 87 hospitalized patients across China and 25 livestock farms. The sequenced isolates were selected from two previous studies (Q. Wang et al., 2018; R. Wang, Liu, et al., 2018). The sequencing reads are available on the Short Read Archive (SRA) under accession number **XXXXXXXX**. All reads were de novo assembled using Unicycler (v0.4.8) (Wick et al., 2017) using default parameters while also specifying hybrid mode for those isolates for which we had both Illumina short-read and PacBio long read sequencing data. Spades (v3.11.1) (Bankevich et al., 2012) was applied, without additional polishing, for cases where Unicycler assemblies failed to resolve. Sequencing datasets without associated metadata on the date of sampling were not included in the analysis.

In total, 2,165 contigs carrying the *bla*_NDM_ gene were identified using BLAST (v2.10.1+) (Camacho et al., 2009). The full metadata table of contigs containing *bla*_NDM_ is available as Supplementary Data 1. The table includes sample accession numbers and information on host organism, collection date, sampling location, assembly status, and contig plasmid type and circularity. Sixteen contigs (C165, C964, GCA_000764615, GCA_000814145, GCA_001860505, GCA_002133365, GCA_002870165, GCA_003194305, GCA_003368345, GCA_003716765, GCA_003860815, GCA_003950255, GCA_003991465, GCA_004795525, GCA_005155965, GCF_004357815) were found to carry more than one copy of *bla*_NDM_ and were not included in our analyses. Two assemblies (GCF_004358085 and GCF_004357805) had a single *bla*_NDM_ gene split into two contigs; these four contigs were also excluded. Contigs GCA_00386065, C184 and C141 were removed due to poor assembly quality. This filtering resulted in a dataset of 2,142 contigs (2,128 genomes) which were used in all subsequent analyses. Of these, six genomes were found to contain *bla*_NDM_ on two contigs, each one harbouring a single copy of *bla*_NDM_.

### Annotating NDM-positive contigs

Coding sequences (CDS) of all NDM-positive contigs were annotated using the Prokka (v1.12) (Seemann, 2014) and Roary (v3.12.0) (Page et al., 2015) pipelines run with default parameters. In addition, plasmid sequences were confirmed based on RefSeq annotation (i.e., contigs labelled “plasmid”), contig circularity reported by Unicycler, or by the presence of a plasmid replicon sequence (Orlek et al., 2017). To identify plasmid replicon types, the contigs were screened against the PlasmidFinder database (version 2020-02-25) (Carattoli et al., 2014) using BLAST (v2.10.1+) (Camacho et al., 2009) where only BLAST hits with a minimum coverage of 80% and percentage identity of >95% were retained. In cases where two or more replicon hits were found at overlapping positions on a contig, the one with the higher percentage identity was retained. All identified plasmid types are provided in Supplementary Data 1.

### Resolving structural variants of NDM-positive contigs

Structural variations upstream and downstream of *bla*_NDM_ were resolved using a novel alignment-based approach, as illustrated in Figure 2. First, contigs carrying *bla*_NDM_ were reoriented such that *bla*_NDM_ gene is on the positive-sense DNA strand (i.e., facing 5’ to 3’ direction). A discontiguous Mega BLAST (v2.10.1+) (Ma et al., 2002) search with default settings was applied against all pairs of retained contigs. This method was selected over the regular Mega BLAST implementation as it is comparably fast, but more permissive towards dissimilar sequences with frequent gaps and mismatches. BLAST hits including a complete *bla*_NDM_ gene on both contigs were selected and cropped to either (i) the start of *bla*_NDM_ gene and the downstream sequence or (ii) the end of the *bla*_NDM_ gene and the upstream sequence depending on the analysis at hand: the downstream or the upstream analysis respectively. This trimming establishes *bla*_NDM_ as an anchor and forces the algorithm to consider only the region upstream or downstream of the gene.

Next, the algorithm proceeds with a stepwise network analysis of BLAST hits. For this purpose, a ‘splitting threshold’ was introduced. Starting from zero, the threshold is gradually increased by 10 bp. At each step, BLAST hits with a length lower than the value given by the ‘splitting threshold’ are excluded. Then, a network is constructed from the remaining BLAST hits such that contigs sharing a BLAST hit are connected with an edge. The network is then broken down into components – groups of nodes (contigs) that share a common edge. It is expected that contigs within each component share a homologous region downstream (or upstream) of *bla*_NDM_ at least of the length given by the threshold. It is therefore not possible for a single contig to be assigned to multiple components. Components of size <5 bp are labelled as ‘Other Structural Variants’ and are not considered in further analyses. Also, contigs that are shorter than the defined ‘splitting threshold’ and share no edge with any other contig are considered as ‘cutting short’.

By tracking the splitting of the network as the ‘splitting threshold’ is increased, one can determine clusters of homologous contigs at any given position downstream or upstream from the anchor gene (here *bla*_NDM_), as well as the homology breakpoint. The precision of the algorithm is directly influenced by the step size which is, in this case, 10 bp and the alignment algorithm, in this case discontiguous Mega BLAST. The described algorithm is available at **LINK**

### Date randomization, linear regression analyses and molecular tip-dating

The 45 complete Tn*125* and 29 complete Tn*3000* contigs harbouring *bla*_NDM_ were sequentially aligned (--pileup flag) using Clustal Omega (v1.2.3) (Sievers et al., 2011) specifying *bla*_NDM_-1 as a profile. The consensus sequence over the alignment was considered the closest match to a putative ancestral sequence and was hence used as a reference to identify SNPs against. This approach was motivated by the fact that: (i) there is no appropriate outgroup sequence available; (ii) the oldest contigs in the dataset can harbour non-ancestral SNPs; (iii) due to a short time span and relatively few mutations present, it is unlikely that any one non-ancestral SNP has become dominant in the population.

Date randomization and linear regression analyses considering the number of SNPs accumulated against the year of sample collection provide an estimate of the strength of the temporal signal in the alignment (Duchene et al., 2019; Rambaut et al., 2016; Rieux & Balloux, 2016). We weighted the linear regressions by the BioProject affiliation of the sequences in the alignments of the two transposons (Supplementary Figure 8A and 8B). This was done to control the strong sampling biases present among samples from the same NCBI BioProject, with contigs from the same BioProject tending to be genetically similar irrespective of the sampling year (Supplementary Figure 9). While both Tn*125* and Tn*3000* showed positive temporal signal (Supplementary Figure 8A and B), neither regression was significant (*p*=0.1279 and *p*=0.1375 respectively). The low sample size and the low genetic diversity in the two alignments may limit the statistical power to detect temporal signal. Date-randomization analysis also showed that the estimated evolutionary rate for both transposons fell within the distribution of slopes on randomized dates (Supplementary Figure 8A and 8B).

A further test of meaningful signal in the data is to consider the degree to which the dated alignment can drive the posterior distribution away from the priors specified in Bayesian dating frameworks. BEAST2 (v2.6.0) (R. Bouckaert et al., 2019) was run on both transposon alignments specifying a strict molecular clock rate with a model averaging prior on the substitution model (R. R. Bouckaert & Drummond, 2017) and a MCMC chain length of 5×10^8^ (Supplementary Data 2). The long MCMC chain length was chosen to ensure convergence. For both runs the Serial Birth-Death Skyline (BDSS) model was specified as the tree prior. The BDSS model is commonly used for viral epidemics (Stadler et al., 2013) which share many parallels with AMR outbreaks. Similar to other birth-death models, the BDSS prior consists of three parameters: a rate of transmission (an estimate transposon/plasmid mobility), recovery (an estimate of transposon fossilization or plasmid loss), and sampling rate. Also, unlike coalescent models, BDSS does not attempt to estimate population sizes, which have limited applicability to dating small genetic regions and mobile elements. We evaluated the prior and posterior distributions across variables after discarding the first 20% of burn-in and after ensuring model convergence (an effective sample size >200).

### Estimating Shannon entropy among NDM-positive contigs

We estimated Shannon entropy (‘diversity’) for several categorizations of *bla*_NDM_-containing contigs: country of sampling, bacterial host genera, replicon type, SNP count within a 5000 bp alignment, and structural variants at positions 3000 bp and 5000 bp downstream of the *bla*_NDM_ gene. The 5000 bp alignment consisted of 654 contigs harbouring *bla*_NDM_, *ble*, *trpF*, *dsbD*, *cutA*, *groS* and *groL* genes. To estimate entropy, we used a weighted bootstrapping approach (1000 iterations) with the probability of pooling any one sample inversely proportional to the number of samples contained in the corresponding BioProject. At each iteration, entropy was estimated for a sampled set of contigs (*X*) classified into *n* unique categories according to the following formula:

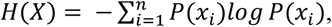

where probability *P(x_i_)* of any sample belonging to any particular category *xi* (e.g., country or replicon type) is approximated using the category’s frequency. Accordingly, higher entropy values indicate an abundance of equally likely categories, while lower entropy indicates a limited number of categories.

### Estimating geographical and genetic distance between contigs

Geographical distance between pairs of selected contigs was determined using the *geodist* (Padgham & Sumner, 2020) R package and reported sampling coordinates or centroids of countries of collection if the former was not available. The exact Jaccard distance (JD) was used as a measure of the genetic distance. It was calculated using the tool Bindash (Zhao, 2019) with *k*-mer size equal to 21 bp. The JD is defined as the fraction of total *k*-mers <u>not</u> shared between two contigs. For instance, JD=1 denotes no *k*-mers are shared. The two distance matrices (genetic and geographic) were assessed using the *mantel* function from *vegan* package in R (Oksanen et al., 2019). To account for the sampling bias, pairs of contigs belonging to the same BioProject were not considered while estimating the Spearman correlation and performing the Mantel test between geographic and genetic distance.

## Acknowledgements

M.A. was supported by a Ph.D. scholarship from University College London. H.W. is supported by National Natural Science Foundation of China (81625014). L.v.D., H.W. and F.B. acknowledge financial support from the Newton Fund UK-China NSFC initiative (MRC Grant MR/P007597/1 and 81661138006). L.v.D. and F.B. are supported from a Wellcome Institutional Strategic Support Fund (ISSF3) – AI in Healthcare (19RX03). F.B. additionally acknowledges support from the BBSRC GCRF scheme and the National Institute for Health Research University College London Hospitals Biomedical Research Centre. L.v.D is supported by a UCL Excellence Fellowship. M.A., L.v.D and F.B. acknowledge UCL Biosciences Big Data equipment grant from BBSRC (BB/R01356X/1). L.P.S. acknowledges funding from the Antimicrobial Resistance Cross-council Initiative supported by the seven UK research councils (NE/N019989/1). The funders had no role in study design, data collection, interpretation of results, or the decision to submit the work for publication. Lastly, M.A. would like to thank Nicola de Maio for informal discussions which led to the idea for the algorithm used to track structural variants.

## Competing interests

The authors declare no financial or non-financial competing interests.

## Contributions

M.A., F.B., L.v.D. and H.W. conceived the project and designed the experiments. M.A., L.v.D., L.P.S., and N.L. collected data from online repositories. R.W., Y.Y., Q.W., S.S, and H.C sequenced samples from Chinese hospitals. M.A., L.v.D, and R.W. *de novo* assembled all the genomes. M.A. performed all the analyses under the guidance of L.v.D and F.B. M.A., L.v.D. and F.B. take responsibility for the accuracy and availability of the results. M.A. wrote the paper with contributions from L.v.D. and F.B. All authors read and commented on successive drafts and all approved the content of the final version.

## Notes

### Competing Interest Statement

The authors have declared no competing interest.

