## Supplementary Material for "Role of the mobilome in the global dissemination of the carbapenem resistance gene *bla*_NDM_"

**Acman et al.**

**Supplementary Table 1. NDM-positive samples stratified by where the data was sourced and the associated sequencing platform.**

| | | Sequencing Platform | | | |
|---|---|---|---|---|---|
| | | ILLUMINA | ION_TORRENT | PACBIO_SMRT & ILLUMINA (Hybrid assembly) | Unknown |
| Source/Database | China Hospitals | 0 | 0 | 112 | 0 |
| | Enterobase | 274 | 0 | 1 | 0 |
| | RefSeq | 0 | 0 | 0 | 1239 |
| | SRA | 515 | 7 | 0 | 0 |

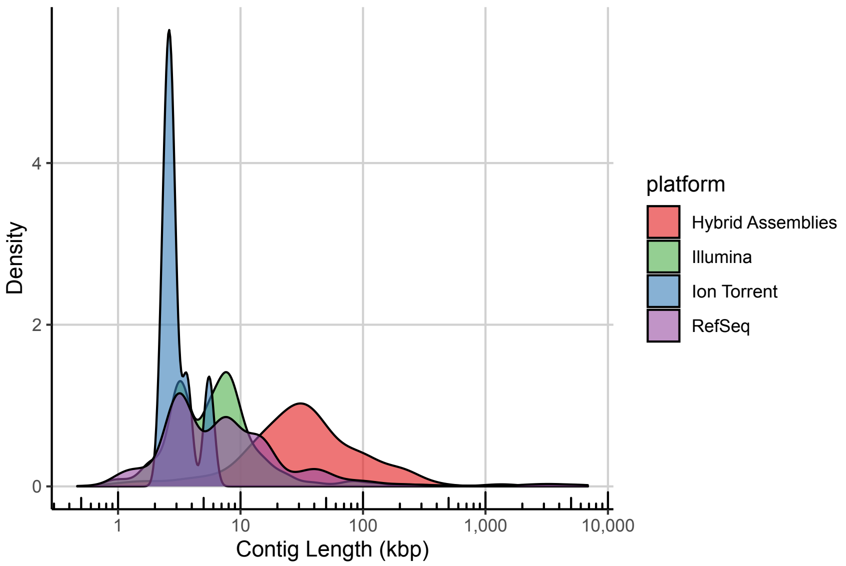


**Supplementary Figure 1. Marginal density distribution of the lengths of all assembled *bla*$_{NDM}$-positive contigs depending on the sequencing platform.**

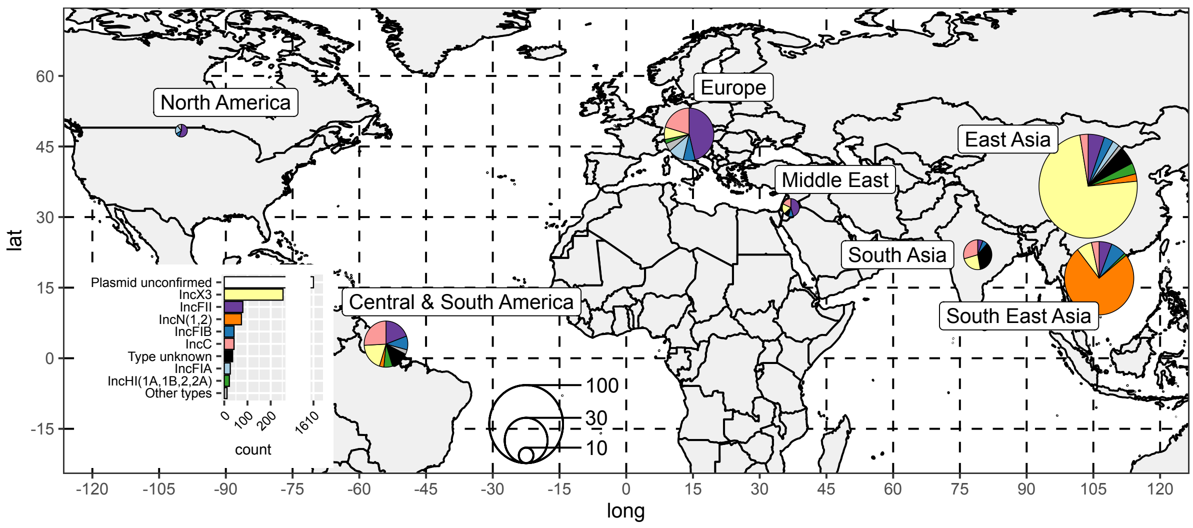


**Supplementary Figure 2. Global distribution of plasmid types of NDM-positive contigs.** All NDM-positive contigs that represent confirmed plasmids (i.e. circularized or of a known plasmid type) have been pooled according to the geographical region and represented on the world map using pie charts. The sizes of the pie charts are log-scaled to aid interpretability.

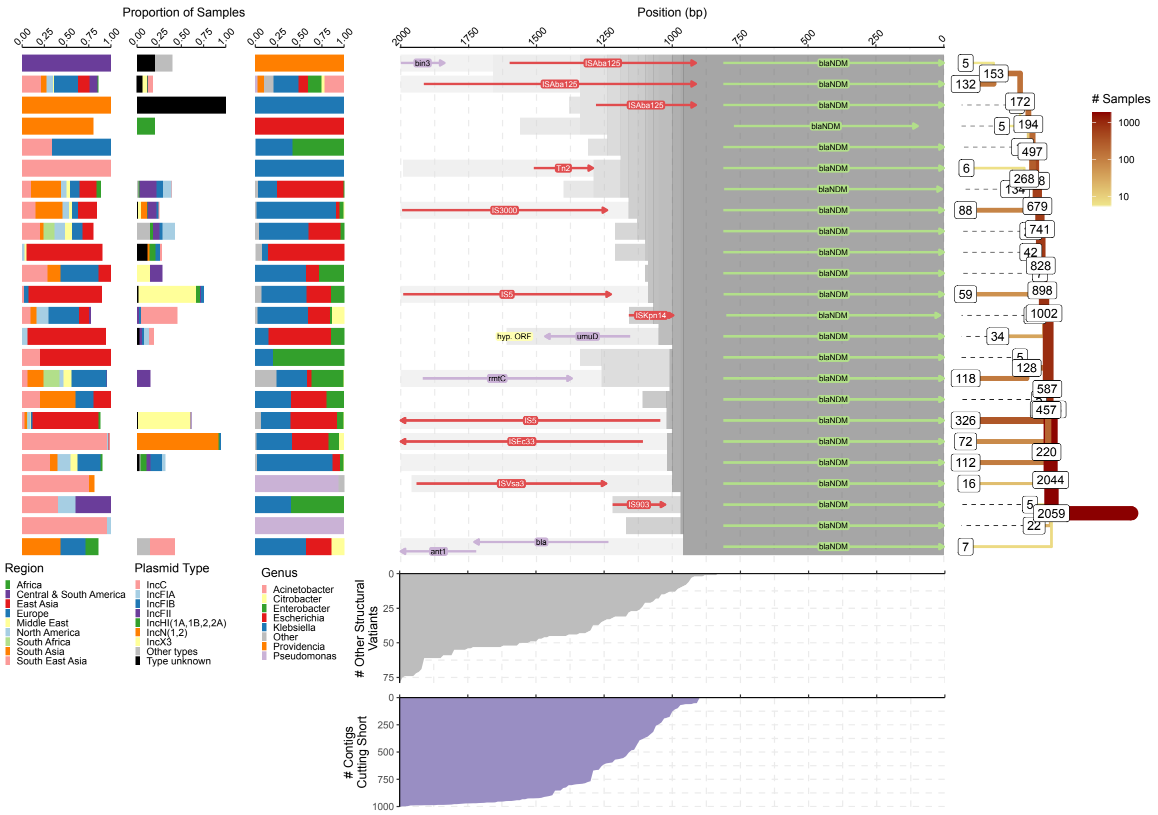


**Supplementary Figure 3. Splitting of structural variants upstream of *bla*$_{NDM}$*.*** The 'splitting tree for the most common (i.e ≥ 5 contigs) structural variants is shown on the right-hand side. The labels on the nodes indicate the number of contigs remaining on each branch. The other contigs either belong to other structural variants or were removed due to being too short in length. The number of contigs cutting short is indicated by the area chart at the bottom. Similarly, the number of contigs belonging to less common structural variants is indicated by the upper area chart. The genome annotations of most common structural variants are shown in the middle of the figure. The homologous regions are indicated by the grey shading. Some of the structural variants and branches were intentionally cut short even though their contigs were of sufficient size. This was done in order to prevent excessive bifurcation and to make the tree easier to interpret. In particular, branches with percent change of contigs lost due to variation and shortness above 30% were truncated. The distribution of genera, plasmid types and geographical regions of samples that belong to a each of the common structural variant is shown on the left-hand side.

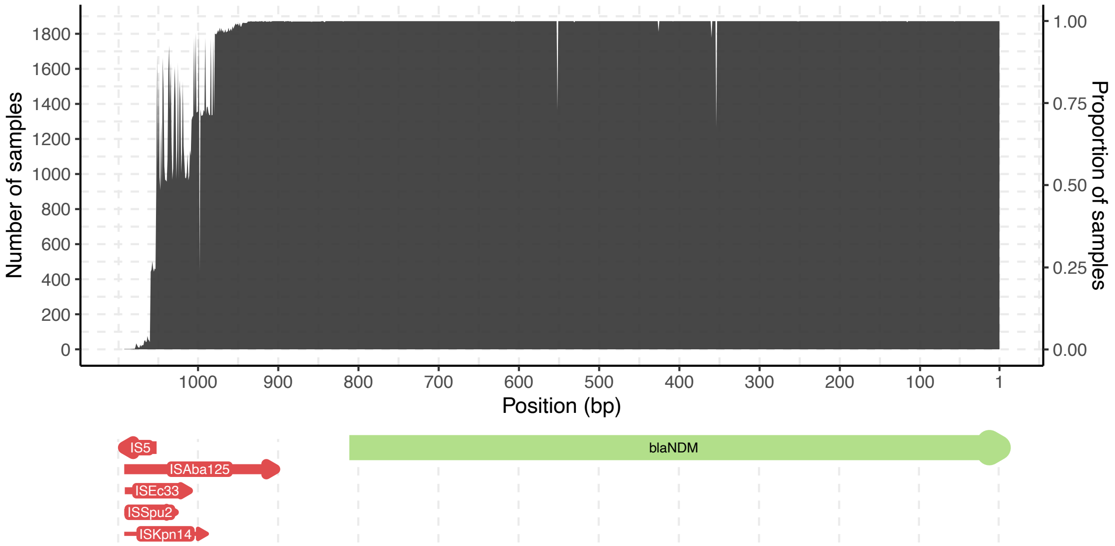


**Supplementary Figure 4. Alignment of 1872 sufficiently long contigs 1050bp upstream of *bla*NDM stop codon.** The alignment also contains 41 gaps mostly at the 3’ end.

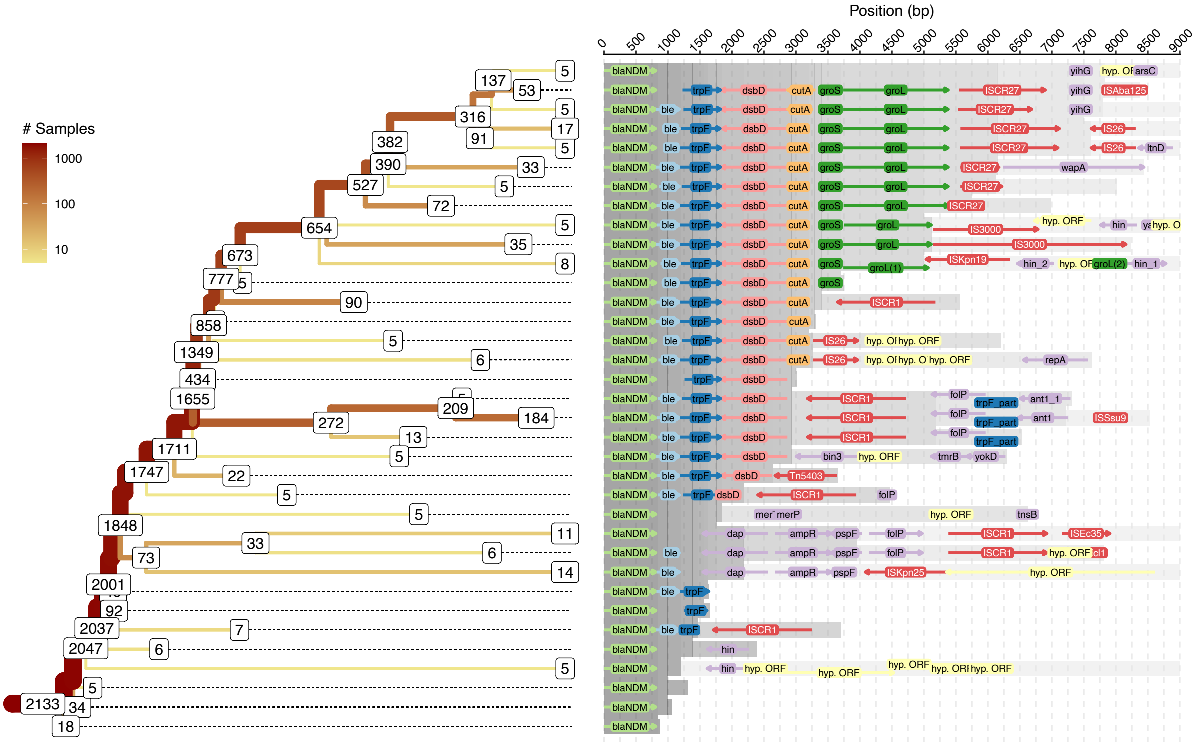


**Supplementary Figure 5. The tree of structural variations downstream of *bla*NDM gene.** This figure is an excerpt from Figure 3 and shows a detailed view of the tree of structural variations and genome annotations of contigs belonging to tree’s terminal branches. The labels on the nodes indicate the number of contigs remaining on each branch. The other contigs either belong to other structural variants or were removed due to being too short in length. The homologous regions are indicated by the grey shading in the genome annotations panel.

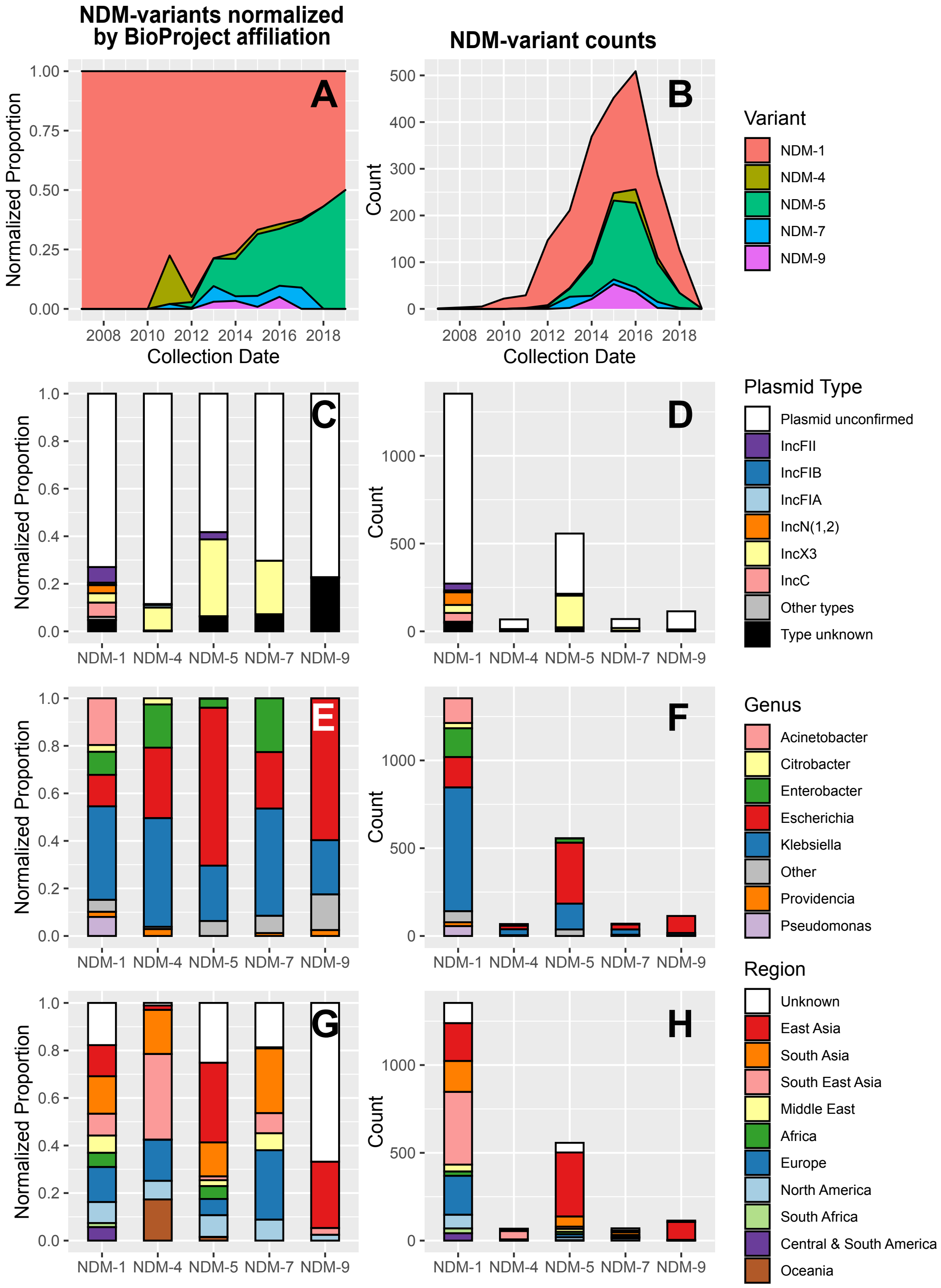


**Supplementary Figure 6. Global prevalence and genetic context of NDM variants.** The left-hand side panels of the figure show the proportions of each category (see legends) normalized by number of BioProjecs within each category while the right-hand side panels show the overall counts. Panels (**A**) and (**B**) show prevalence of the five most common (i.e., >10 contigs) NDM variants over time. The remaining panels show bar plots indicating normalized proportions and counts respectively of plasmid types (**C** and **D**), bacterial isolate genus (**E** and **F**) and sampling geographical location (**G** and **H**) across five NDM variants.

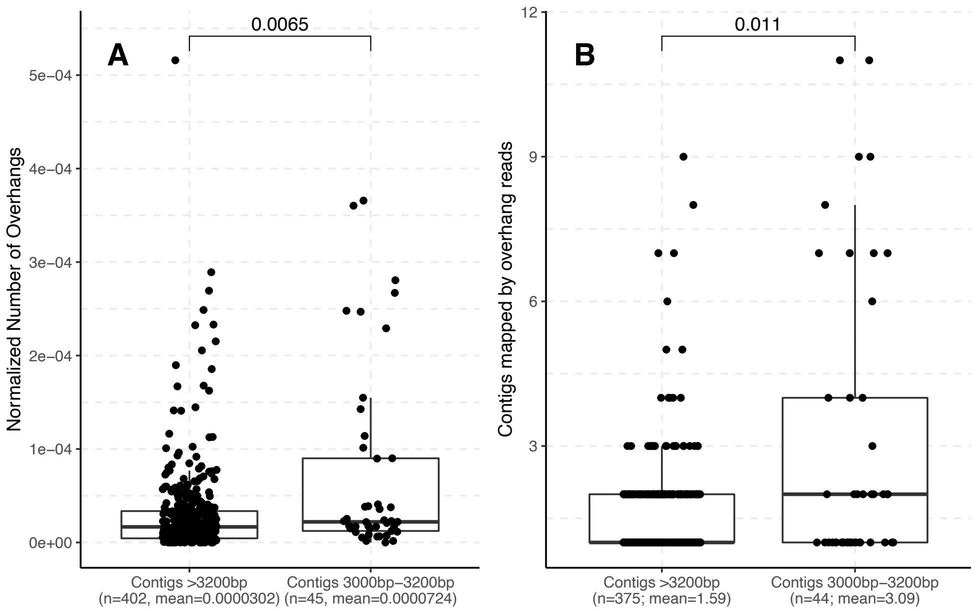


**Supplementary Figure 7. Overhanging reads of contigs truncated between 3000-3200bp downstream of *bla*NDM are on average more abundant (A) and map to multiple contigs (B).** Four hundred sixty-seven short-read sequencing samples from SRA have been used for this analysis. The reads overhanging the right (3') flank of contigs carrying *bla*NDM have been identified and counted. The abundance of overhanging reads was normalized by the total number of sequencing reads in the corresponding sample and then compared between two groups of contigs using Mann–Whitney U test (**A**). Next, the overhanging reads ≥50bp were mapped back to the contigs of the corresponding sample and the number of mapped contigs was counted and compared between the two groups (**B**).

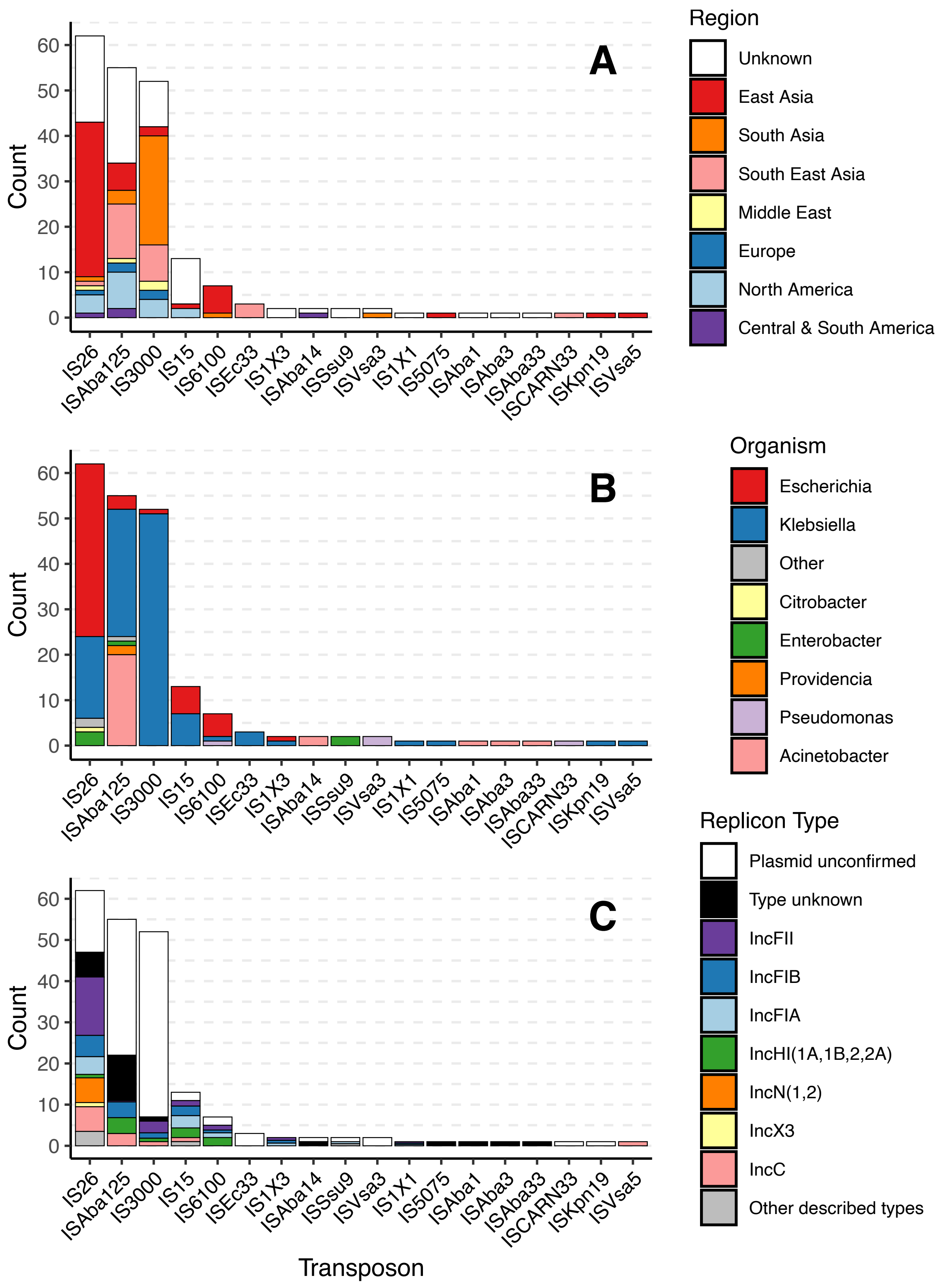


**Supplementary Figure 8. Barplots showing the number of contigs within the dataset with two copies of the same insertion sequence (ISs) less than 30kbp apart that surround the *bla*$_{NDM}$ gene.** The two copies of the same IS are presumed to form a composite transposon. The bar plots in the figure are the same and they have been coloured by geographical region (**A**), bacterial host genera (**B**) and contig replicon type (**C**).

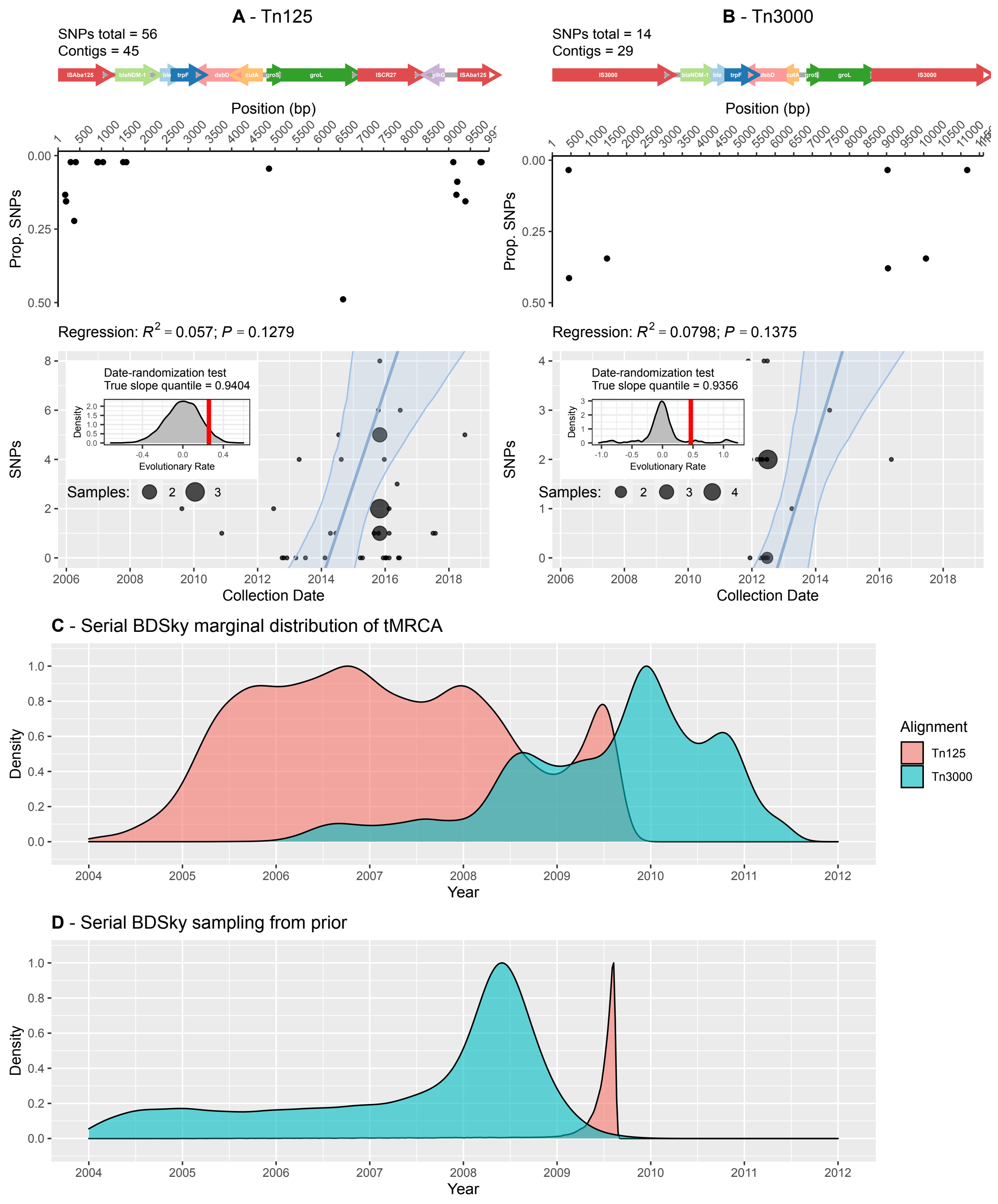


**Supplementary Figure 9. Results of the date randomization and linear regression analyses (A, B) and Bayesian molecular tip-dating (C, D) of the alignments of Tn125 and Tn3000 composite transposons carrying *bla*$_{\mathbf{NDM}}$ gene.** Starting from the top, both panels **A** and **B** contain the following plots: schematic representation of the aligned transposon sequences, the SNP frequency across the alignment, and the weighted linear regression analysis of number of SNPs accumulated against the year of sample collection. To account for the sampling bias in the dataset (Supplementary Figure 9), linear regression was weighted based on the sample BioProject affiliation. Specifically, each data point in the regression was given a weight equal to 1/*n* where *n* is the corresponding BioProject size. The ribbon surrounding the regression line provides a 95% confidence interval given by the bootstrapping the regression analysis (1000 iterations). The inset plot shows the results of the date-randomization test. The marginal distribution of the inset indicates the regression line slope values (i.e., evolutionary rates) after 10,000 date randomizations and the red vertical line indicates the true slope value. The BEAST2 runs were performed with the following priors: strict clock rate, (Supplementary Data 2), substitution model averaging, and a Serial Birth-Death Skyline (Serial BDSky) phylogenetic tree prior. The resulting marginal distributions of the most recent common ancestor (tMRCA) estimates for Tn125 and Tn3000 are shown in panel **C**. The panel **D**, on the other hand, shows the BEAST2 tMRCA estimates after sampling from priors (i.e., no SNP data provided to the model)

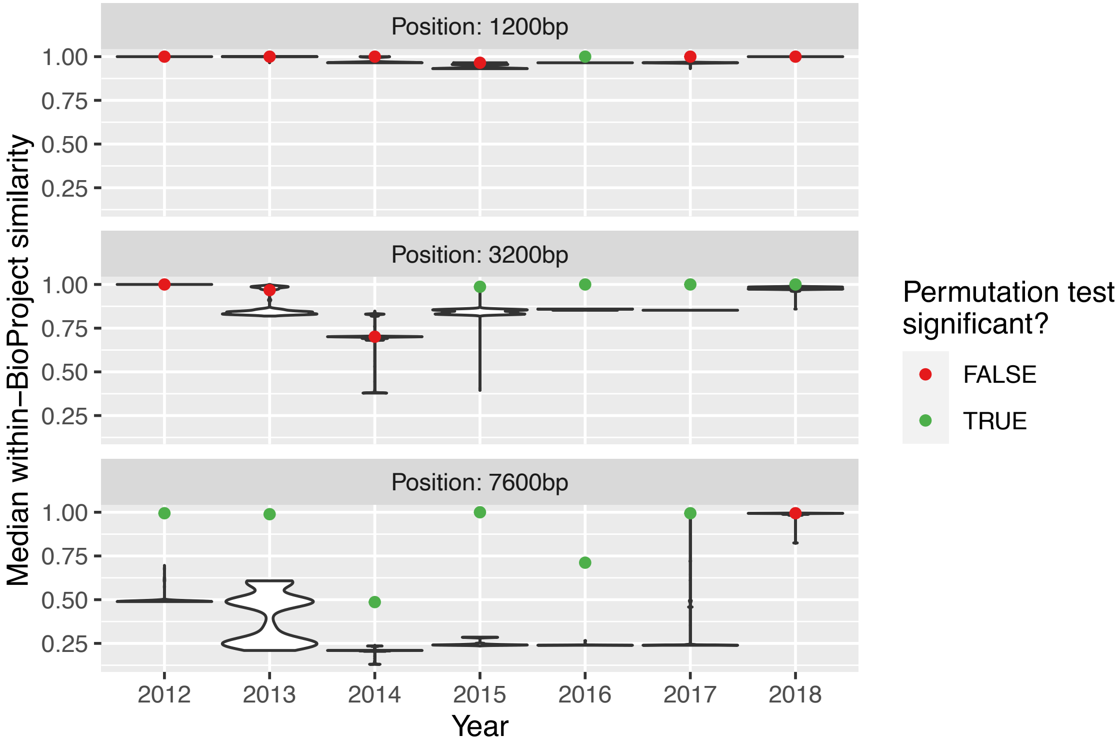


**Supplementary Figure 10. Permutation test reveals strong within-BioProject sampling bias.** Permutation tests were performed on contigs from BioProjects of size > 2 over a range of collection years (2012-2018) and for three positions downstream of $bla_{\mathrm{NDM}}$ (1200bp, 3200bp and 7600bp). Years with less than 5 BioProjects with more than 2 samples were not considered. Sequence similarity between all pairs of contigs was estimated using the minhash algorithm. The median within- BioProject sequence similarity is indicated by points. The violin plots indicate the within-BioProject sequence similarity after randomization of BioProject labelling (1000 iterations). The colouring of the points indicates if the true median within-BioProject similarity falls outside 95% of randomization interval (<2.5$^{th}$ or >97.5$^{th}$ percentile).

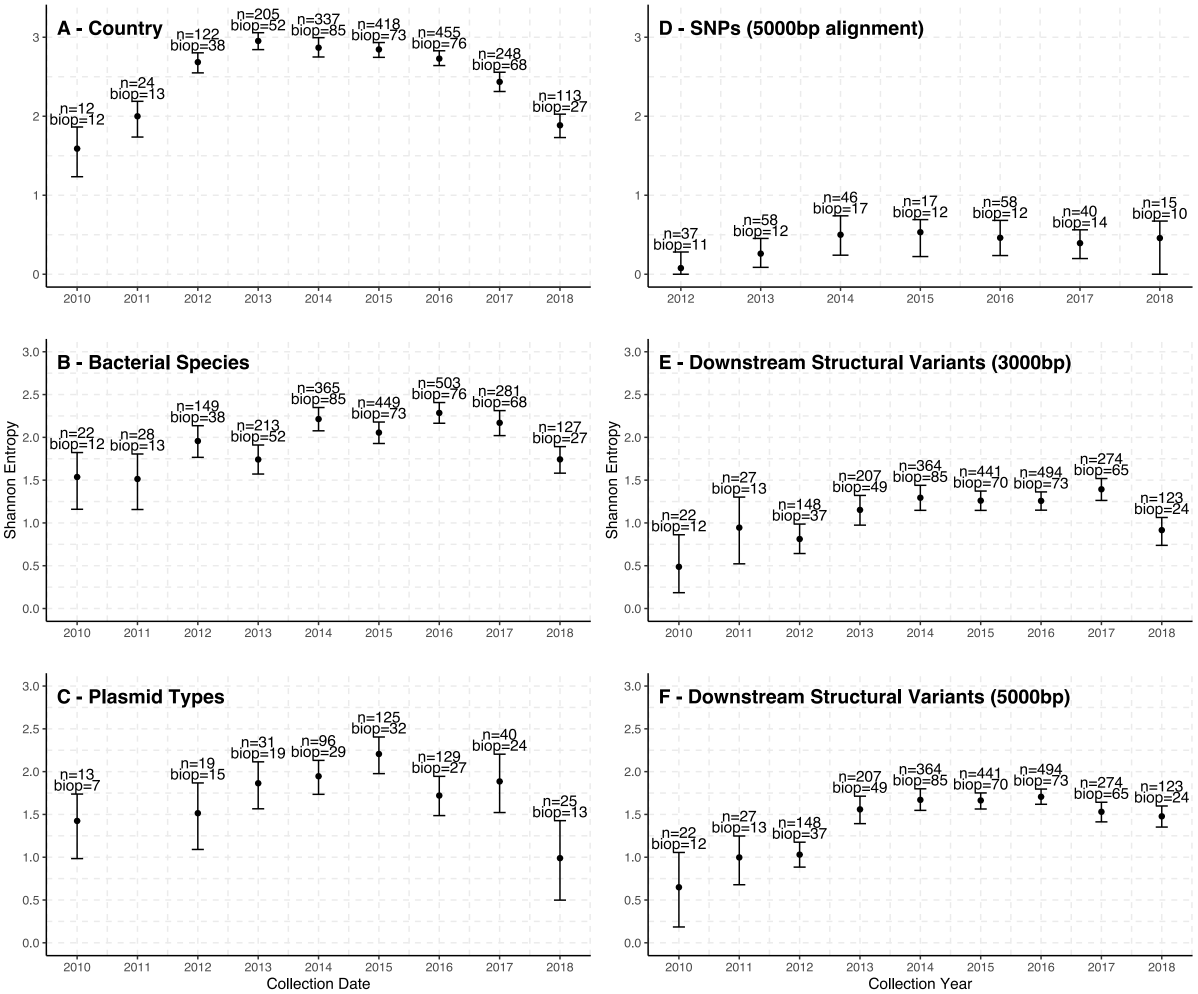


**Supplementary Figure 11. Change in Shannon entropy (diversity) over time for six categories of NDM-positive samples.** More specifically, the entropy was estimated for samples' country labelling (**A**), bacterial genera (**B**), replicon types of contigs bearing *bla*NDM gene (**C**), SNP counts within 5000bp alignment of *bla*NDM*-ble-trpF-dsbD-cutA-groS-groL* genes downstream of blaNDM (**D**), structural variants 3000bp (**E**) and 5000bp downstream of *bla*NDM (**F**). The median entropy (points) together with 95% confidence interval (error bars) was estimated using weighted bootstrapping (1,000 iterations). In particular, sample selection while bootstrapping was adjusted based on BioProject affiliation to account for the sampling bias in the dataset (Supplementary Figure 10). This was done in similar fashion to the linear regression analysis (Supplementary Figure 9A and B). Each sample had a 1/n probability of being selected by the bootstrapping algorithm, where *n* corresponds to the BioProject size. Years with too few samples (≤10) were excluded from the analysis.

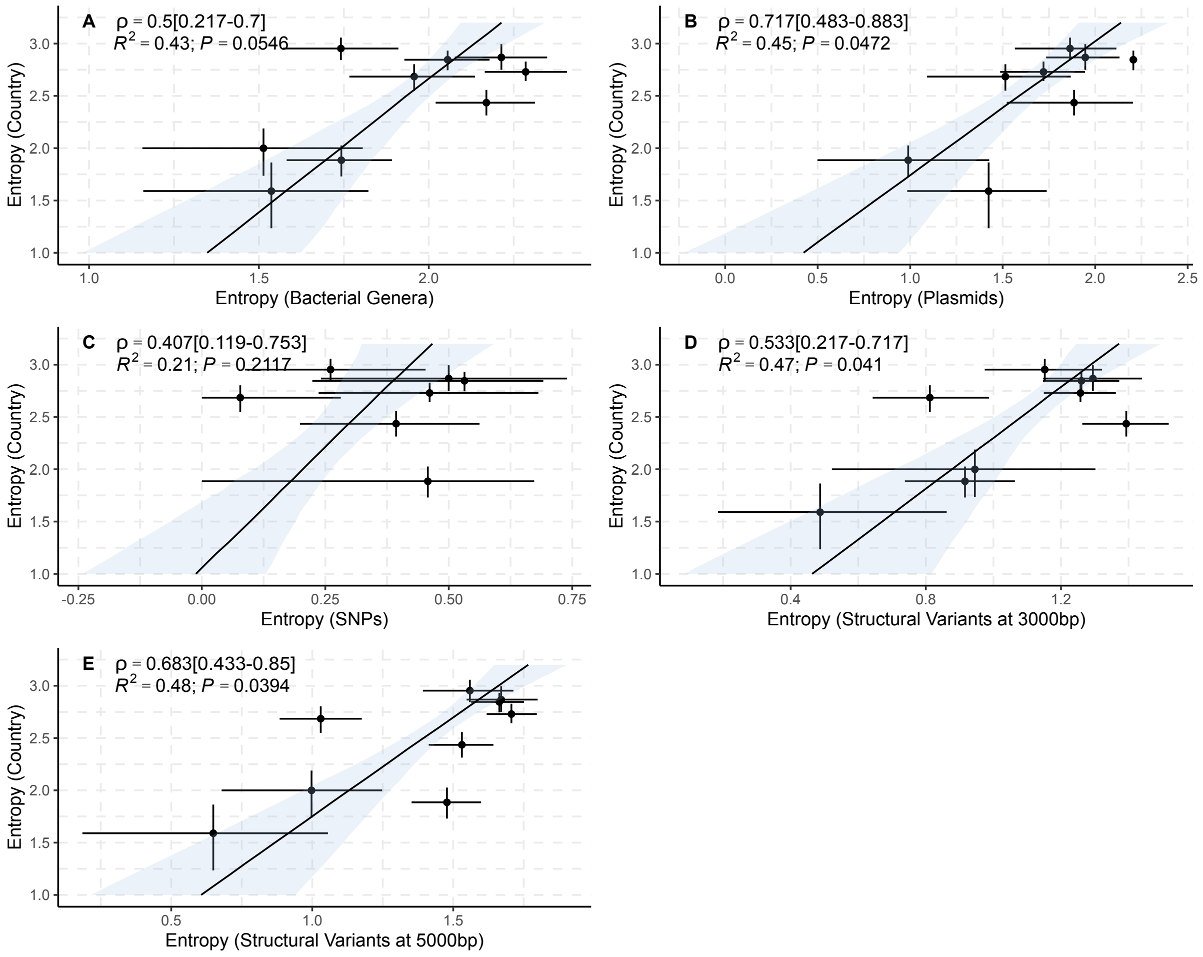


**Supplementary Figure 12. Spearman correlation and linear regression of Shannon entropy (diversity) estimates.** The Shannon entropy bootstrapped values (Supplementary Figure 11) were used to provide a median and 95% confidence interval (CI) of Spearman correlation coefficients, as well as a median regression line with 95% CI (ribbon) between samples' country labelling and: bacterial genera (**A**), replicon types (**B**), SNP counts within 5000bp alignment downstream of *bla*NDM (**C**), and structural variants 3000bp (**D**) and 5000bp downstream of *bla*NDM (**E**).

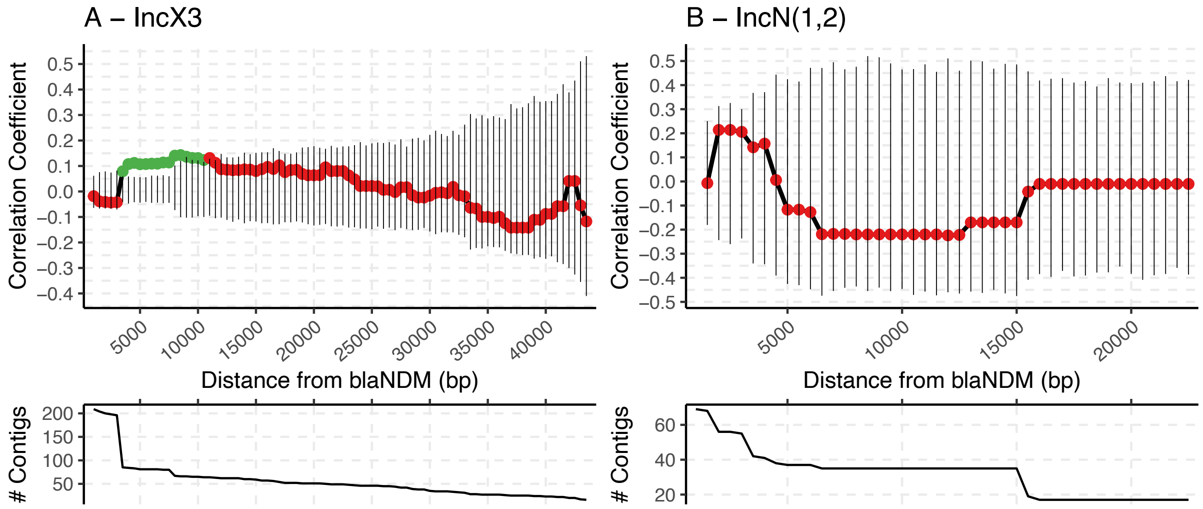


**Supplementary Figure 13. A graph indicating the Spearman correlation estimates between genetic and geographical distance of IncX3 and IncN plasmids as the DNA sequence upon which the genetic distance is measured is increased downstream of *bla*NDM gene.** Same as in Figure 4, the exact Jaccard index was used as a measure of genetic distance and was determined using alignment-free BinDash tool (v0.2.1) and the geographical distance was estimated with *geodist* (v0.0.6) R package using sampling coordinates or sampling country centroids if the former had not been provided. The analysis was performed on two groups of contigs: the ones carrying *bla*NDM gene on a confirmed IncX3 plasmid (**A**) and the ones carrying the gene on confirmed IncN1 or IncN2 plasmids (**B**)**.** In both cases, the genetic and geographical distance was measured between all pairs of contigs from a different BioProject which yielded two distance matrices: genetic and geographical. The Spearman correlation was then estimated between the two matrices and its significance evaluated using Mantel (randomization) test. The significant Spearman correlation (p-value <0.05) was indicated with the green point and the non-significant one with the red point, while the black vertical lines indicate the 95% confidence interval of 1000 Mantel test permutations. The genetic distance matrix and subsequent Spearman correlation were estimated multiple times by increasing the assessed DNA sequence starting from *bla*NDM gene and continuing downstream. The two plots below the correlation graphs indicate the number of contigs used in the correlation analysis as the assessed DNA sequence is increased.
